# Sex differences in early human fetal brain development

**DOI:** 10.1101/2024.03.04.583285

**Authors:** Federica Buonocore, Jenifer P Suntharalingham, Olumide K Ogunbiyi, Aragorn Jones, Nadjeda Moreno, Paola Niola, Tony Brooks, Nita Solanky, Mehul T. Dattani, Ignacio del Valle, John C. Achermann

## Abstract

The influence of sex chromosomes and sex hormones on early human brain development is poorly understood. We therefore undertook transcriptomic analysis of 46,XY and 46,XX human brain cortex samples (n=64) at four different time points between 7.5 and 17 weeks post conception (wpc), in two independent studies. This developmental period encompasses the onset of testicular testosterone secretion in the 46,XY fetus (8wpc). Differences in sex chromosome gene expression included X-inactivation genes (*XIST*, *TSIX*) in 46,XX samples; core Y chromosome genes (n=18) in 46,XY samples; and two Y chromosome brain specific genes, *PCDH11Y* and *RP11-424G14.1*. *PCDH11Y* (protocadherin11 Y-linked) regulates excitatory neurons; this gene is unique to humans and is implicated in language development. *RP11-424G14.1* is a novel long non-coding RNA. Fewer differences in sex hormone pathway-related genes were seen. The androgen receptor (*AR*, NR4A2) showed cortex expression in both sexes, which decreased with age. Global cortical sex hormone effects were not seen, but more localized AR mechanisms may be important with time (e.g., hypothalamus). Taken together, our data suggest that limited but potentially important sex differences occur during early human fetal brain development.

## Introduction

The human brain undergoes remarkable growth and differentiation during the first and second trimester^1,2^, but many of the mechanisms that influence these processes remain poorly understood. In recent years, rapid progress has been made in the development of techniques that can analyze gene and protein expression in the brain across the lifespan, and an increasing number of resources are becoming available to share data and promote scientific progress^3–6^. One major area of interest has been the potential impact of biological sex differences on neurobiological development and function^7^. However, in humans, most reported studies in this area have focused on post-natal or adult life, and surprisingly few data are available related to sex differences during critical early stages of brain development^8^.

Early developmental sex differences can potentially result from the influences of *sex chromosomes* as well as the effects of sexually dimorphic *sex hormones* in early life. A complex interplay between these elements and other factors may exist^7,9^.

The sex chromosome complement is established at the time of fertilization as a result of sex chromosome combination. Females typically have a 46,XX karyotype and males typically have a 46,XY karyotype, although biological variation does occur^10^. The X chromosome encodes more than 800 genes and has a small number of genes with Y chromosome homologues in the pseudoautosomal regions. In contrast, the Y chromosome encodes for only around 60 genes that play a key role in sex development and fertility, as well as more diverse biological functions^11^.

Sex hormones are generally synthesized and released by the developing gonads (testes, ovaries). In the 46,XY embryo, the Y chromosome gene *SRY* is expressed in the bipotential gonad at around 6 weeks post conception (wpc), which triggers a cascade of downstream genes leading to testis determination^12–14^. We have previously shown by modeling time-series analysis of gene expression that maximal upregulation of the enzymes needed to synthesize testosterone occurs at 8wpc in the human testis^12^. Fetal testicular testosterone acts to stabilize Wolffian structures (e.g., seminal vesicles, vas deferens) and is converted to the more potent hormone, dihydrotestosterone (DHT) by the enzyme 5α-reductase type 2 (*SRD5A2*) in the external genital region to promote penile and scrotal growth^15^. Testosterone acts exclusively through the androgen receptor (*AR*) (also known as *NR3C4*) and can be converted to estrogens by the enzyme aromatase (*CYP19A1*). Most data suggest that the developing ovary in the 46,XX fetus is endocrinologically quiescent at this time and does not secrete significant amounts of the female-typical hormone, estradiol.

Sex chromosomes and sex hormones play important dynamic roles during development, leading to longer term effects after birth, at puberty and in adult life, for example, shaping the way that specific conditions may affect males and females differently. These affect not just reproductive and endocrine conditions, but also influence many biological processes such as cardiovascular, immune, and neurological function. For example, in the field of clinical neuroscience, autistic spectrum disorder (ASD), conduct disorders, schizophrenia and Parkinson’s disease are more common in males, whereas anxiolytic disorders and dementia are typically more prevalent in females^16^. Sex differences can therefore potentially play a central role both in disease mechanisms and in the way specific conditions manifest.

In order to address sex differences in early human brain cortex development further, we undertook transcriptomic analysis across a critical time period of development between 7.5-8wpc and 15-17wpc, at a time just before the onset of fetal testicular testosterone secretion and in subsequent weeks. The main aims of the study were to address whether: 1) global differences exist between 46,XX and 46,XY samples that could represent the effect of sex chromosome-related genes, and whether this differs in the brain compared to other tissues; 2) whether any differentially expressed genes have strong brain specificity; 3) whether sex-related divergent patterns in global cortical gene transcription occur over time and whether these could potentially represent sex-hormone (testosterone)-dependent events; and 4) whether more localized sex hormone effects might occur in the developing brain cortex or other regions that could be linked to biological outcomes.

## Results

### Global sex differences in gene transcription during early human brain cortex development

In order to investigate potential *global* sex differences in gene transcription in brain development, 32 samples were obtained from developing human brain telencephalon/cortex between Carnegie Stage 22-23 (CS22-CS23), corresponding to 7.5-8wpc, and 15-17wpc (termed “Brain-Seq 1”) (Figure 1a). An overview of the study design is shown in Table 1, which included matched 46,XX and 46,XY samples at each of four different stages (CS22-23, 9wpc, 11-12wpc, 15-17wpc). This time period represents a key phase in brain cortex development and differentiation, and follows the onset of testosterone secretion by the 46,XY testis from around 8wpc (Figure 1b & c).

**Figure 1.**
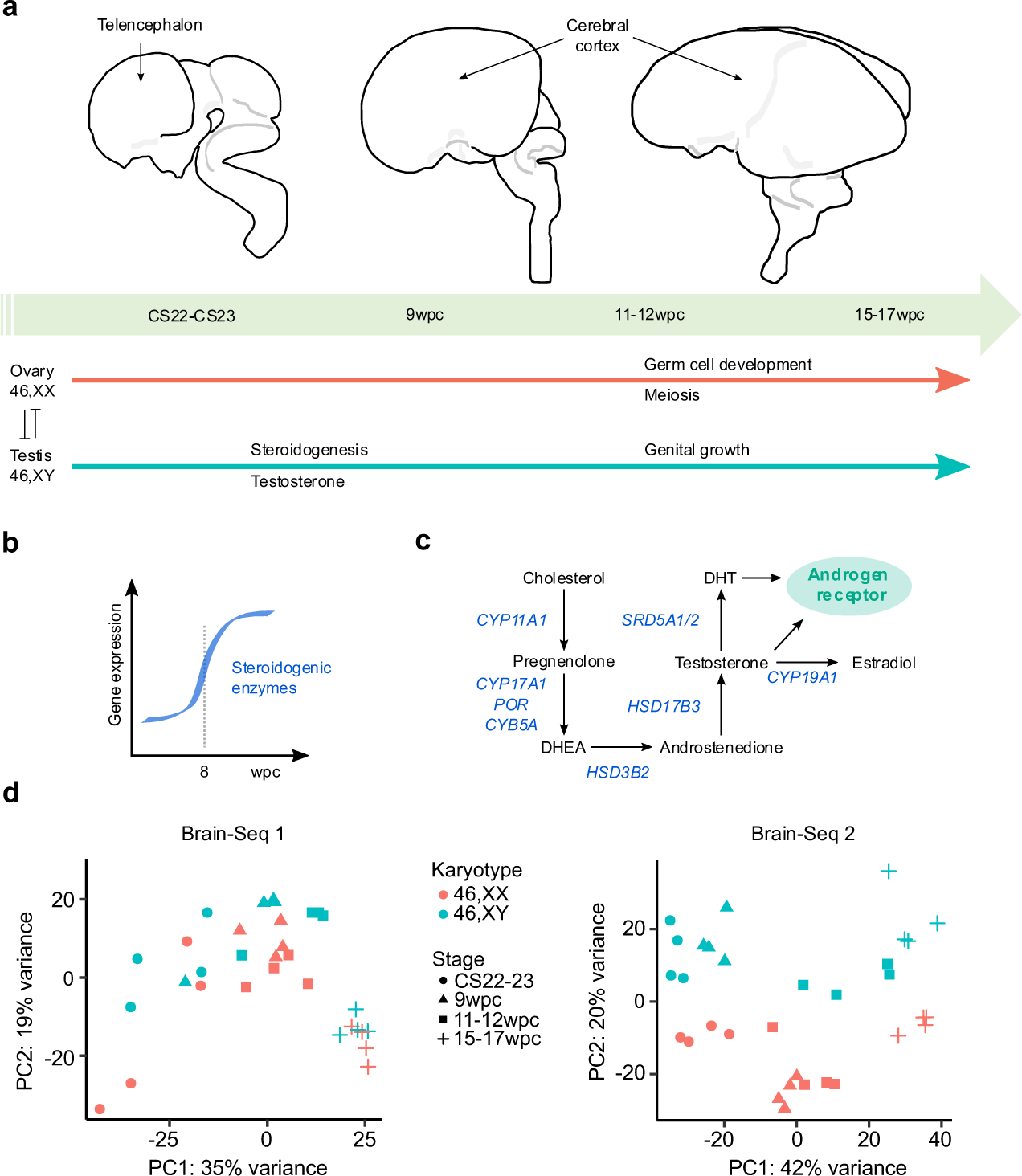
Global sex differences during early brain development. **a** Model of human brain development in relation to gonad development. Telencephalon and cerebral cortex regions of the developing brain are indicated by arrows. Gonad determination into either testis or ovary begins at around Carnegie Stage 18 (CS18) (6 weeks post conception, wpc). Testicular testosterone synthesis and secretion occurs from around CS23 (8wpc) in the 46,XY fetus. **b** Gene expression of factors implicated in testicular testosterone biosynthesis is upregulated at 8wpc. Data derived from Del Valle et., al^12^. **c** Simplified pathway of androgen biosynthesis with genes encoding enzymes showed in blue. DHEA, dehydroepiandrosterone; DHT, dihydrotestosterone. **d** Principal component analysis (PCA) for each independent bulk RNA-seq dataset (Brain-Seq 1, Brain-Seq 2) included in the study showing all samples (total n=32 in each dataset, see Table 1) based on the first two principal components (PC1, PC2).

**Table 1.** Overview of all samples used in the study. Description of brain cortex samples analyzed in our study, showing the number of 46,XX and 46,XY samples per each developmental stage. CS22 corresponds to 7wpc+4d(days); CS23 corresponds to 8wpc. n, number; wpc, weeks post-conception. Major groups are shown in bold; sub-groups contributing to these numbers are shown below them in non-bold,

|  | Brain-Seq 1 dataset |  |  | Brain-Seq 2 dataset |  |  |
| --- | --- | --- | --- | --- | --- | --- |
| Developmental stage | 46,XX (n) | 46,XY (n) | Total (n) | 46,XX (n) | 46,XY (n) | Total (n) |
| **CS22-23** | **4** | **4** | **8** | **4** | **4** | **8** |
| CS22 | 4 | 4 | - | - | 3 | - |
| CS23 | - | - | - | 4 | 1 | - |
| **9wpc** | **4** | **4** | **8** | **4** | **4** | **8** |
| **11-12wpc** | **4** | **4** | **8** | **4** | **4** | **8** |
| 11wpc | 4 | 4 | - | 4 | - | - |
| 12wpc | - | - | - | - | 4 | - |
| **15-17wpc** | **4** | **4** | **8** | **4** | **4** | **8** |
| 15wpc | 3 | 3 | 8 | - | - | - |
| 16wpc | 1 | 1 | - | 4 | - | - |
| 17wpc | - | - | - | - | 4 | - |
| **Overall** |  |  | **32** |  |  | **32** |

**Table 2.**
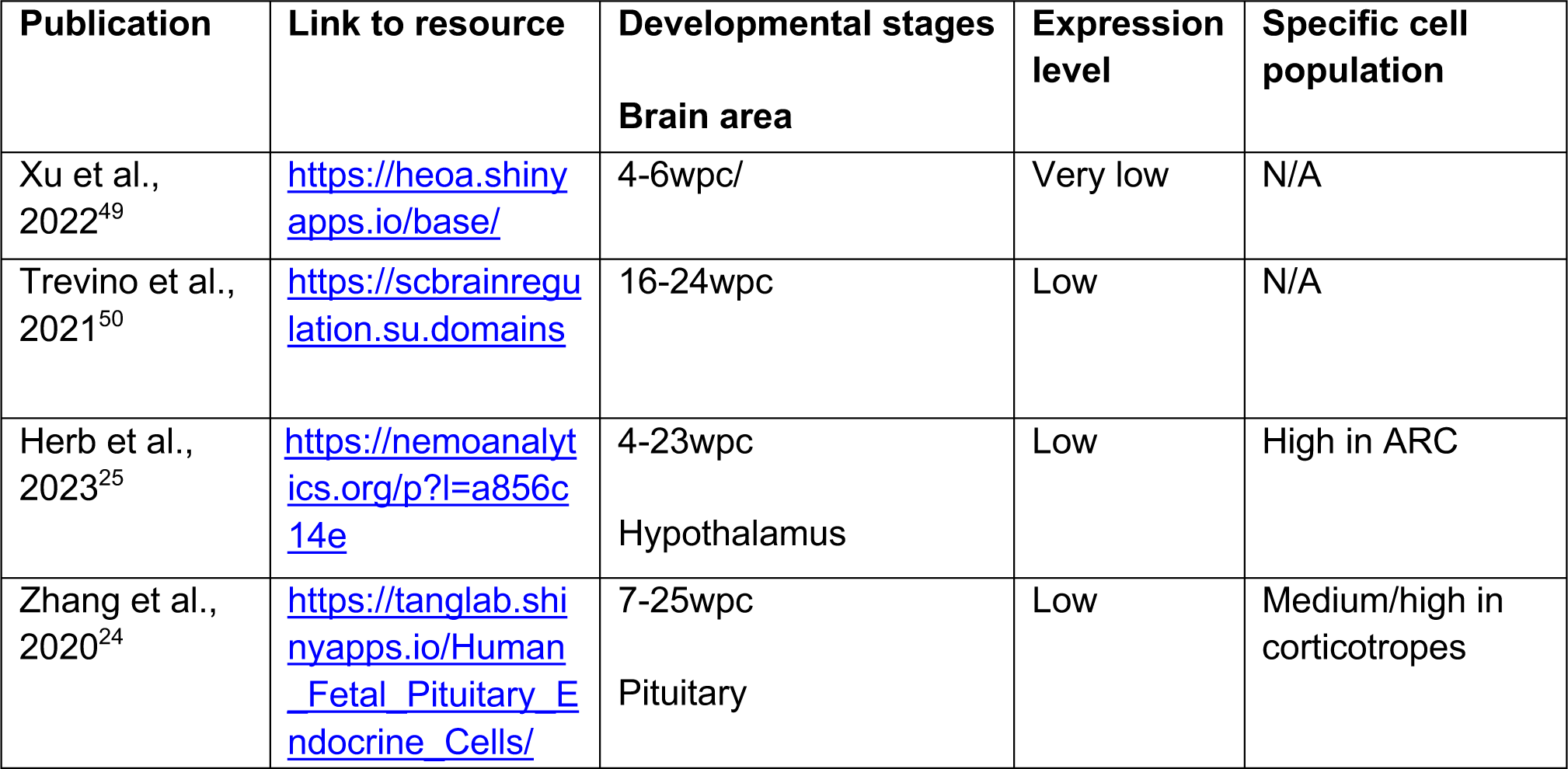
Overview of AR expression in published single cell RNA-Seq studies of human brain development. Data were taken from publicly available resources. ARC, arcuate nucleus. N/A, not available.

A matched bulk RNA-Seq dataset was also generated using the same bioinformatic pipeline from raw data available as part of a Human Developmental Biology Resource (HDBR) fetal brain transcription repository (Table 1)^17^. This dataset allowed a parallel, independent replication and validating study across the same developmental time course (termed “Brain-Seq 2”).

Principal component analyses (PCA) for both Brain-Seq 1 and Brain-Seq 2 datasets are shown in Figure 1d. Principle component (PC) 1 clearly reflected developmental stage of the sample (Brain-Seq 1, 35% variance; Brain-Seq 2, 42% variance), whereas the second component (PC2) likely reflected karyotype of the sample and any potential sex differences in transcription (Brain-Seq 1, 19% variance; Brain-Seq 2, 20% variance).

Differential expression of key sex chromosome-related genes was confirmed in volcano plots of 46,XX versus 46,XY samples at each stage and in both Brain-Seq 1 and Brain-Seq 2 datasets (Figure 2). Using a cut-off of log2 fold change (FC) >0.7 and adjusted p-value of <0.05, only two genes showed consistently higher expression in all 46,XX samples; these were the two key regulators of X inactivation, *XIST* and *TSIX* (Figure 3a, 3c, 3e). In contrast, a “core” group of 18 genes were consistently highly expressed in 46,XY brain samples, which were all Y chromosome genes (Figure 3b, 3d, 3f). However, analysis of bulk-RNA-seq from other development tissues (e.g. kidney, pancreas, liver, skin) also showed a component of dimensionality related to karyotype, suggesting that at least some non-specific differences in sex chromosome gene transcription occur in development (Supplementary Figure 1).

**Figure 2.**
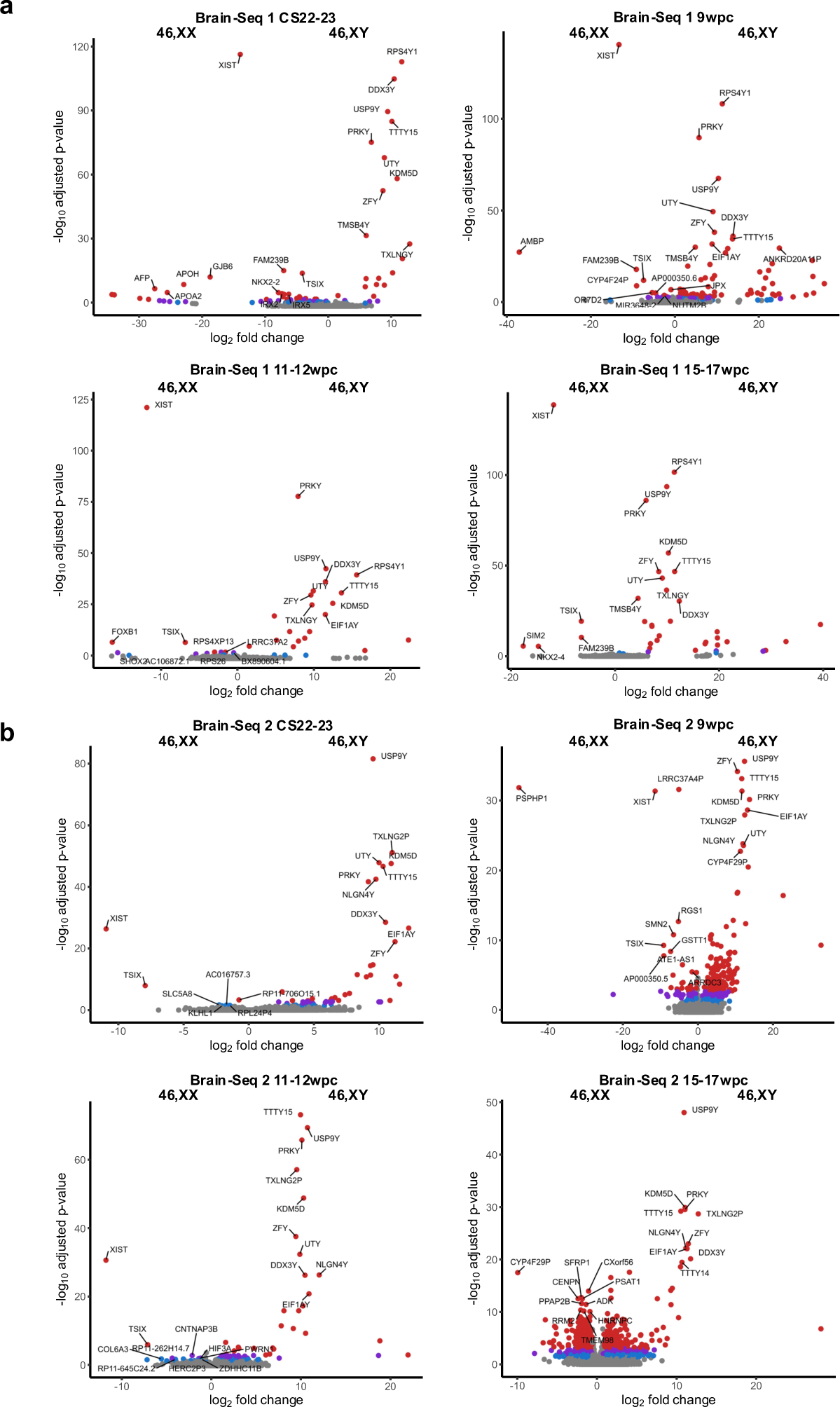
Volcano plots for differentially expressed genes between 46,XX and 46,XY samples at each developmental stage. **a** Brain-Seq 1 dataset. **b** Brain-Seq 2 dataset. At each developmental stage there are n=4 samples in the 46,XX group and n=4 samples in the 46,XY group. Red indicates adjusted p-value <0.001; purple indicates adjusted p-value <0.01; blue indicates adjusted p-value <0.05. The ten genes with the highest −log10 adjusted p-value in each group are labeled (where adjusted p-value <0.001). CS, Carnegie stage; wpc, weeks post conception.

**Figure 3.**
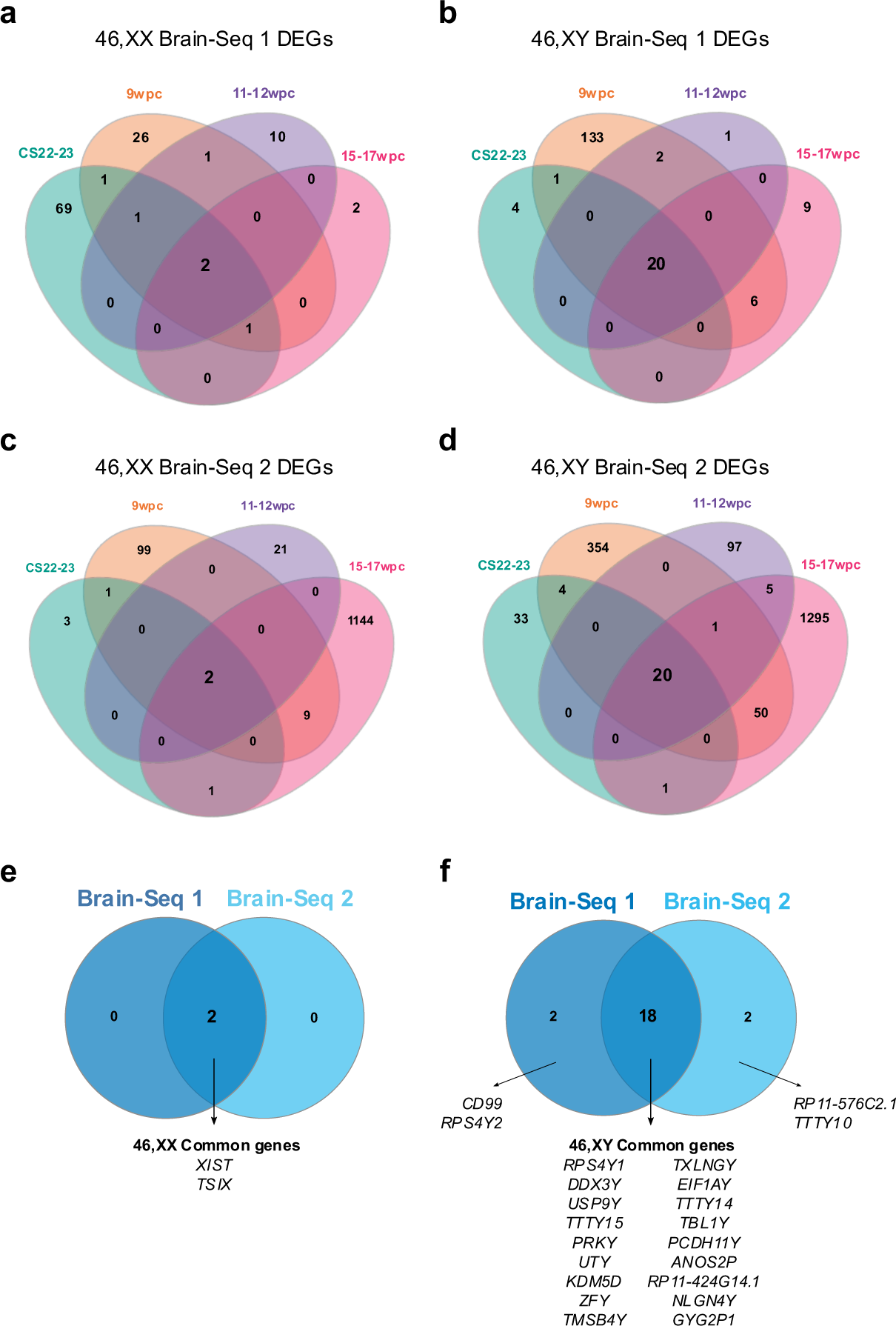
Sex-specific differential gene expression patterns during different stages of human brain development. **a-d** Venn diagrams showing the overlap of differentially expressed genes (DEGs) in both RNA-seq datasets at each developmental stage (CS22-23; 9wpc; 11-12wpc; 15-17wpc). **a** 46,XX Brain-Seq 1 dataset; **b** 46,XY Brain-Seq 1 dataset; **c** 46,XX Brain-Seq 2 dataset; **d** 46,XY Brain-Seq 2 dataset. Differential expression was defined as a log2 fold change >0.7 and adjusted p-value <0.05. Those genes found to be shared across all four stages in each dataset were identified and then compared between the two datasets. **e** Common 46,XX differentially expressed genes in both datasets. **f** Common 46,XY differentially expressed genes in both datasets. CS, Carnegie stage; wpc, weeks post conception.

### Brain-specific sex differences in gene transcription

In order to identify *brain-specific* genes during development, differentially expressed 46,XY genes (46,XY versus 46,XX, log2FC >0.7, adjusted p-value <0.05) in the brain cortex (Figure 3f) were compared with differentially expressed 46,XY genes in the other tissues (kidney, pancreas, liver, skin) (Figure 4a). Using this approach only two genes emerged as showing strong brain specific expression (*PCDH11Y*, *RP11-424G14.1*). *PCDH11Y* encodes protocadherin 11 Y linked, a cadherin-family extracellular adhesion molecule involved in cell-cell communication in excitatory neurons (https://www.proteinatlas.org/ENSG00000099715-PCDH11Y). Significantly higher expression was confirmed in the developing 46,XY brain compared to 46,XX brain and to 46,XY control tissues by more detailed analysis of the bulk RNA-seq data and by qRT-PCR (Figure 4b, 4c, 4d). These time series-analyses of the bulk RNA-seq datasets and qRT-PCR revealed a marked increase in *PCDH11Y* in the 46,XY brain with age, especially at 15-17wpc (Figure 4b, 4d). Consistent with this is the differential expression of *PCDH11Y* in the adult brain, compared to other tissues, in Human Protein Atlas Consensus and Genotype-Tissue Expression (GTEx) data (Figure 4e, Supplementary Figure 2).

**Figure 4.**
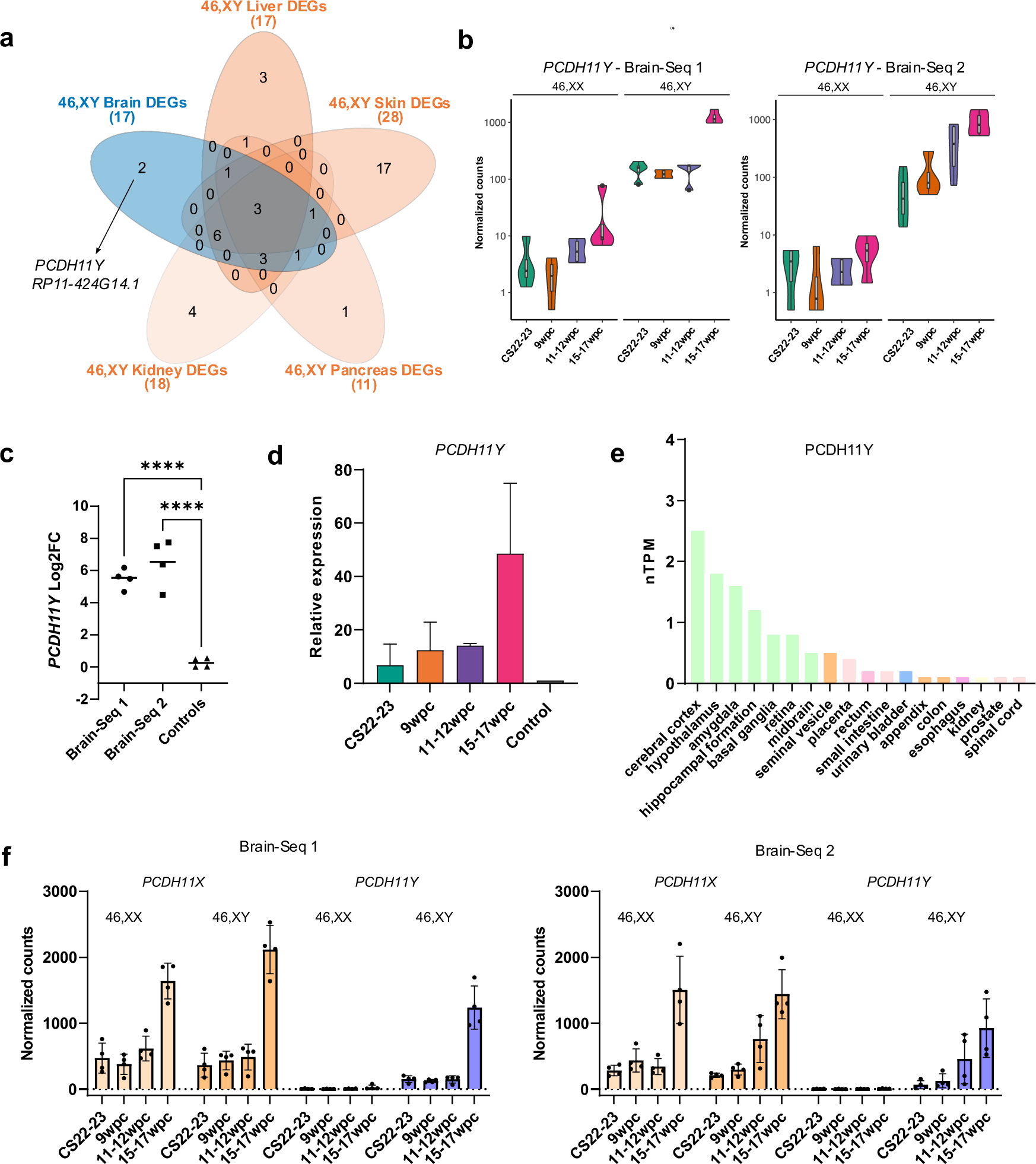
Analyses of brain-specific genes in 46,XY cortex. **a** Venn diagram showing the overlap of 46,XY differentially expressed genes (DEGs) common to both brain datasets and several 46,XY independent control tissues. In this analysis, *TTTY15* was not included as it mapped to USP9Y in the control datasets. **b** Violin plots of normalized counts showing expression patterns of *PCDH11Y* across developmental stages in the 46,XX and 46,XY brain and in both datasets (n=4 in each stage).**c** Log2 fold change (Log2FC) values of *PCDH11Y* in both brain datasets compared to controls across each developmental stage; horizontal line represents the median value. Statistical analysis was performed using a one-way ANOVA with Dunnett’s multiple comparison tests; ****p-value <0.001. **d** qRT-PCR analysis of *PCDH11Y* in 46,XY brain cortex across key developmental stages, and compared to control. Two separate experiments were performed; in each, three independent samples were used for each stage, each done in triplicate. Four control tissues were used (kidney, liver, pancreas, skin). Representative data from one experiment is shown as mean with standard deviation. **e** Expression of *PCDH11Y* in the Human Protein Atlas Consensus dataset for adult brain, showing highest expression in cerebral cortex and hypothalamus, as well as in other brain structures (data accessed and downloaded from https://www.proteinatlas.org). nTPM, normalized transcript per million. **f** Relative expression of *PCDH11X* and *PCDH11Y* in the Brain-Seq 1 and Brain-Seq 2 datasets (n=4 in each group); data are shown as mean with standard deviation. CS, Carnegie stage; wpc, weeks post conception.

*PCDH11Y* is located on the short arm of the Y chromosome (p11.2) (Chr Y:5,000,226-5,742,224, GRCh38) in a region with strong X chromosome homology. Although *PCDH11Y* is unique to humans, a related X chromosome gene (*PCDH11X*) exists (Supplementary Figure 3). In both our Brain-Seq 1 and Brain-Seq 2 datasets, *PCDH11X* expression was similar in both 46,XX and 46,XY samples (Figure 4f). *PCDH11X* was seen in the brain and some other tissues in GTEx, with no clear sex differences (Supplementary Figure 4). Thus, the Y chromosome gene *PCDH11Y* likely has an additive or unique effect in the developing and post-natal 46,XY brain.

The other brain-specific differentially expressed transcript identified using this analysis pipeline was a long non-coding RNA, *RP11-42G14.1* (also referred to as ENSG00000260197) (Yq11.222) (Chr Y:19,691,941-19,694,606, GRCh38.p14) (Supplementary Figure 5). The gene with closest proximity to *RP11-424G14.1* is *KDM5D* (Chr Y:19,703,865-19,744,939), but both have a minus strand orientation and *RP11-424G14.1* is 3’ to *KDM5D* (Supplementary Figure 5). Little is known about the putative function of this gene.

### Brain-enriched Y chromosome genes

In addition to “brain-specific” genes identified using this approach, we also investigated the potential differential expression of “brain-enriched” genes from the Y chromosome. These genes demonstrated differential expression in the developing 46,XY brain but also more variable degrees of differential expression in different control tissues (46,XY versus 46,XX), meaning they did not reach the threshold for detection in our pipeline based on log2FC, adjusted p-value, or both.

The main genes identified were *ANOS2P, DDX3Y, EIF1AY, GYG2P1, KDM5D, NLGN4Y, PRKY, RSP4Y1, TBL1Y, TMSB4Y, TTTY14, TXLNGY, USP9Y, UTY, and ZFY*, with remarkable consistency between both Brain-Seq1 and Brain-Seq2 datasets. As expected, these genes fall within the “core” set of 18 genes and are represented within the context of other Y chromosome genes in Supplementary Figure 6. We therefore explored expression of these genes across tissues in more detail using the Human Protein Atlas Consensus RNA expression panel of 50 different tissues (including 10 neuronal-related regions), but no genes with high brain specificity were identified, although *NLGN4Y* shows brain expression as well as expression in multiple other tissues (Supplementary Table 1). *PCDH11Y* emerged as the Y gene with highest brain specificity.

### Sex hormone effects on global early human brain development

In addition to differences in sex chromosome-related transcripts, differences in sex hormones and their pathways are also likely to influence sex dimorphic aspects of brain development during gestation.

As outlined above, the 46,XY human embryo/fetus develops testes that start to synthesize and release androgens (testosterone) into the developing blood stream from around 8wpc (Figure 1b)^12^. Testosterone typically acts through the androgen receptor (*AR*, *NR3C4*), and in some tissues such as the developing genital tubercle/external genitalia, testosterone has to be converted to the more potent androgen, dihydrotestosterone (DHT) (Figure 1c). The role of estrogens (e.g., estradiol) in the developing brain is unclear, although the ovary is not thought to synthesize significant amounts of estrogen during early development (Figure 1c).

To address potential sex hormone pathways in the developing brain, we first analyzed *AR* gene expression across time in the Brain-Seq 1 and Brain-Seq 2 fetal brain cortex datasets. Both approaches showed a remarkably similar level of *AR* expression in 46,XY and 46,XX tissues, and a consistent and marked decrease in *AR* expression with age (Figure 5a, 5b). This finding was consistent with generally low levels of AR expression in the adult brain, and globally similar AR expression in males (46,XY) and females (46,XX) in GTEx (Supplementary Figure 7). Furthermore, the number of differentially expressed genes in the 46,XY cortex over time was similar to the 46,XX cortex, suggesting that a clear and obvious prolonged divergence of potential androgen-driven target genes did not occur (Supplementary Figure 8). In support of this, additional time series analyses of DEGs (log2 fold change >0.7 and p-adj <0.05) that are higher at 9wpc compared to CS22-23, as well as higher at 15-17wpc compared to CS22-23 in 46,XY samples in both brain datasets, showed no consistently up regulated androgen responsive genes (Supplementary Figure 9).

**Figure 5.**
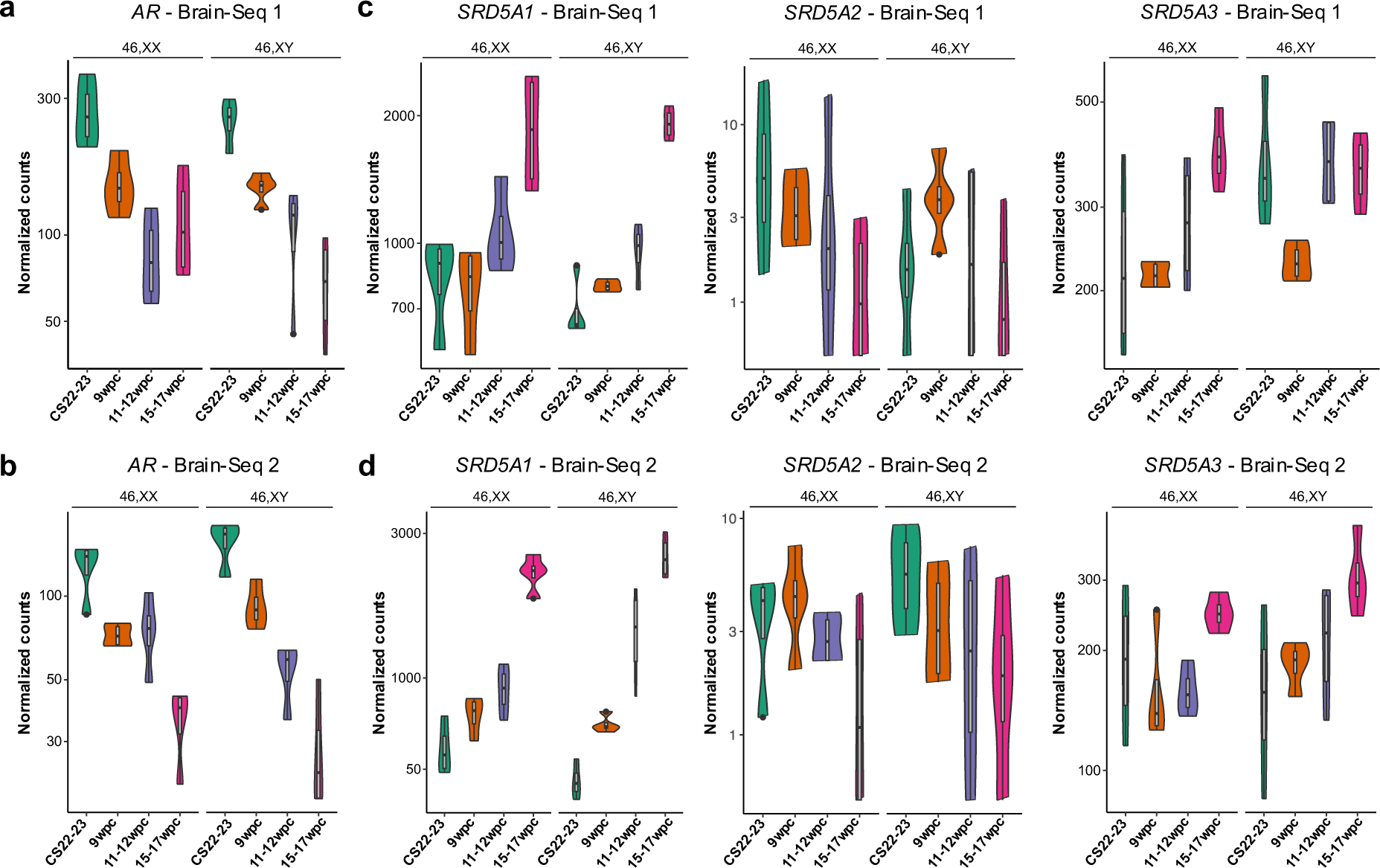
Expression of genes encoding the androgen receptor (*AR*) and related factors involved in dihydrotestosterone biosynthesis. **a** Violin plots of normalized counts showing expression patterns of the *AR* in the Brain-Seq 1 dataset (n=4 in each group) and **b** Brain-Seq 2 dataset. **c** Violin plots of normalized counts showing expression patterns of the *SRD5A1, SRD5A2* and *SRD5A3* enzyme-encoding genes in the Brain-Seq 1 dataset (n=4 in each group) and **d** Brain-Seq 2 dataset. CS, Carnegie stage; wpc, weeks post conception.

By comparing 46,XX and 46,XY DEGs of both Brain-Seq1 and Brain-Seq2 datasets with a database of AR interacting proteins (Reactome KnowledgeBase)^18^, several factors were differentially expressed in the developing brain (Supplementary Figure 10), although overall there was no consistent enrichment of AR interactors in the dataset.

We also investigated the brain cortex expression of enzymes involved in “classic” and “non-classic/backdoor” pathways for DHT generation^19–21^ (Supplementary Figures 11 & 12). Very low expression of *SRD5A2* was seen, the enzyme in the classic pathway that generally coverts testosterone to DHT in reproductive tissues (e.g. developing penis, prostate) (Figures 1c, 5c and d). *SRD5A1* and *SRD5A3* enzyme isoforms were expressed with similar levels in 46,XY and 46,XX cortex (Figure 5c and d). The expression of other proteins and enzymes involved in the “classic” and “backdoor” pathways of androgen biosynthesis were extremely variable in the brain cortex, although an almost complete lack of *CYP17A1* expression (encoding 17α-hydroxylase/17,20-lyase) would likely be rate limiting; for example, in the potential neurosteroidogenesis of androgens from precursors such as placental progesterone or fetal adrenal gland androsterone^21–23^ (Supplementary Figures 11 & 12).

As estrogens are proposed to play a key role in brain sex differences in some species, we investigated these pathways further in our human fetal datasets. Expression of the gene encoding aromatase (*CYP19A1*) (needed to convert androgens to estrogens, Figure 1c) and of the estrogen receptors (*ESR1*, *ESR2*) all had extremely low counts in all cortex samples and did not show any sex dimorphic differences (Supplementary Figure 13).

Taken together, these data show that the androgen receptor gene (*AR*) is expressed in early human fetal brain cortex in both 46,XY and 46,XX fetuses, and that there is a marked decrease in *AR* expression in both sexes in the weeks following the onset of testicular testosterone release, but many alternative pathways linked to sex steroid generation and action are not strongly expressed. Finally, no potentially androgen responsive genes showed specific and consistent upregulation in the 46,XY brain cortex samples.

### Potential localized androgen receptor expression

Given the detectable *AR* expression seen at a transcript level in both 46,XX and 46,XY brain cortex, we undertook immunohistochemistry using an AR specific antibody that was validated in human development. Although most areas of cortex did not show strong androgen receptor expression, clear regions of AR positivity were seen in the nuclei of cortical cells in early development (Figure 6a). Furthermore, more localized regions of AR expression were seen including in midline structures and the basal hypothalamus (Figure 6b and Supplementary Figure 14 and 15).

**Figure 6.**
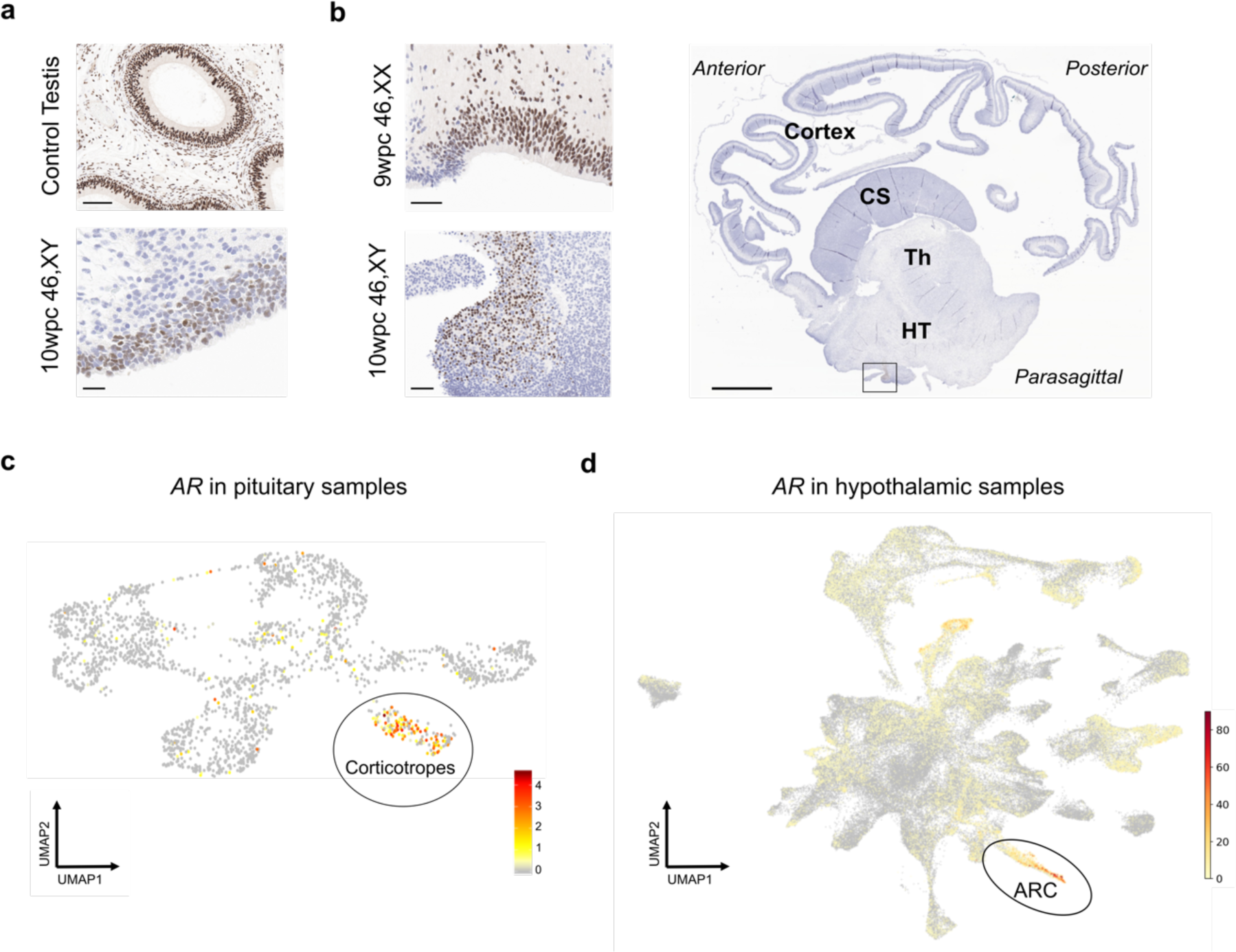
Androgen receptor expression in the developing human brain. **a** Immunohistochemistry (IHC) of control prepubertal testis showing nuclear androgen receptor expression (upper panel; scale bar 100 µm) and 10 week post conception (wpc) cerebral cortex (46,XY) (lower panel; scale bar 20 µm). **b** IHC of localized regions of AR expression in the 9 wpc cortex (46,XX) and in a hypothalamic region at 10 wpc (46,XY) (scale bars 50 µm). The orientation of this localized AR staining in the 10wpc brain is indicated by the rectangle in a parasagittal section (right panel). CS, corpus striatum/ganglionic eminence; HT, hypothalamus; Th, thalamus. **c** Single cell RNA-seq expression of androgen receptor (*AR*) in human pituitary samples (7-25wpc) (total n=21; 46,XX n=11; 46,XY n=10). High expression of AR is shown in red in pituitary corticotropes (Supplementary Figure 15). Data accessed from https://tanglab.shinyapps.io/Human_Fetal_Pituitary_Endocrine_Cells/ (CC BY 4.0, https://creativecommons.org/licenses/by/4.0/)^24^. **d** Single cell RNA-seq expression of androgen receptor (*AR*) in human hypothalamus samples (4-23wpc) (total n=11; 46,XX n=4; 46,XY n=7). High expression of AR is shown in red in the arcuate nucleus (ARC) of the hypothalamus (Supplementary Figure 15). Data accessed from https://nemoanalytics.org/p?l=a856c14e (CC BY-NC 4.0, https://creativecommons.org/licenses/by-nc/4.0/)^25^. UMAP, uniform manifold approximation and projection.

In order to address this further, we analyzed several available resources that investigated transcriptomic expression during early human embryonic and fetal brain development (Table 2). Using the GTEx dataset in adult tissues across the lifespan, highest expression of *AR* was seen in the anterior pituitary and hypothalamus regions (Supplementary Figure 7; see also https://www.proteinatlas.org/ENSG00000169083-AR/brain). We therefore focused on these regions in more detail, as they are also involved in hypothalamic-pituitary-endocrine regulation. Using a human fetal pituitary gland single cell RNA-sequencing (scRNA-seq) analysis resource, we identified highest AR expression in the corticotrope cell lineage, suggesting a potential role in the adrenal axis and later stress mechanisms^24^ (Figure 6c, Table 2). Furthermore, in a human fetal hypothalamus single cell dataset we were able to localize AR expression in several areas, most strongly in the arcuate nucleus (ARC)^25^ (Figure 6d, Table 2). Taken together, these findings suggest that localized expression of the *AR* may be important with time.

## Discussion

Although sex differences in early development are potentially biologically important, relatively little is known about these at a genetic level in the late first trimester and early second trimester in humans. We therefore undertook this time-series study looking at potential effects of sex chromosomes and the machinery for sex hormone action across this time frame in the developing human cerebral cortex.

We intentionally chose to focus on a large, well-matched bulk RNA sequencing approach. Whilst this does not provide the granular detail of transcriptomics in single cells, it has the advantage of allowing detection of low-level gene expression and trends, especially in time series data, and has less sampling bias than might occur with a more focal single cell approach. In order to strengthen the robustness of this strategy, we included two matched replication datasets of 32 samples each at similar time points (Brain-Seq 1 and Brain-Seq 2), which were balanced for 46,XY and 46,XX samples. We also chose a study period spanning the developmental stages just before testicular testosterone secretion in the 46,XY fetus, and in the weeks after its onset, in order to assess any basal sex differences and any potential divergent changes in gene expression which could be due to differences in sex hormone action.

Our analysis initially focused on *global* differences in gene expression across this critical time period. Using principal components analysis, a strong effect of developmental stage (age) was seen (PC1). An effect of karyotype at each stage was generally seen in PC2, and more pronounced in the Brain-Seq2 dataset. A noticeable divergence of “clustering” in the weeks following the onset of testosterone secretion (8wpc) in males was *not* obvious, suggesting most sex-dependent transcriptomic activity during this study period up until 15-17 wpc is relatively fixed and predominantly reflects sex chromosome gene expression rather than major differences in sex hormone action at a global level. This hypothesis is supported by our finding that all consistently differentially expressed genes (46,XX>46,XY and 46,XY>46 XX) across the time course were encoded by the sex chromosomes: that is, X chromosome genes involved in X inactivation (*XIST*, *TSIX*) in 46,XX samples, and a core group of 18 Y chromosome genes in 46,XY samples. No autosomal gene differences were consistently seen at the cut-off thresholds used and during the developmental stages studied, although we cannot exclude that these could be seen with time. Furthermore, when a supplementary analysis of sex differences in development was performed in other tissues (kidney, pancreas, liver, skin), somewhat similar patterns of differences in the principal components for 46,XX and 46,XY was seen. The differences in each of these tissues also mostly represented core X chromosome and Y chromosome genes. Taken together, sex differences in early brain (cortex) gene expression were seen, but these largely reflected sex chromosome gene effects during this developmental time period.

In order to identify potential *brain-specific* genes, we intersected our differentially expressed brain genes with those differentially expressed in other tissues. The two X chromosome genes involved in X inactivation (*XIST*, *TSIX*) were clearly differentially expressed in all 46,XX tissues, as expected, and were not brain specific. Of the 18 Y chromosome genes identified, only two emerged as highly brain specific; *PCDH11Y* and the non-coding *RP11-424G14.1* (ENSG00000260197). Of note, several other Y chromosome genes showed strong brain expression in both datasets, but also had variable expression in other tissues. These genes could also have important developmental effects in the brain and in different organs, but also did not in general show strong brain specificity in adult transcriptomic datasets. Therefore, for the purposes of this study, we chose to focus mainly on the likely brain-specific genes.

The most relevant brain-specific Y chromosome gene identified was *PCDH11Y*. *PCDH11Y* encodes “protocadherin 11 Y-linked” (OMIM 400022). This protein is a proposed transmembrane adhesion molecule thought to be involved in neural development and synaptic cell-cell communication. *PCDH11Y* is expressed in adult brain datasets, and single cell analysis suggests that the transcript clusters in neuronal excitatory/inhibitory synaptic neurons (Human Protein Atlas). *PCDH11Y* is located on the short arm of the Y chromosome (p11.2) in a region with strong X chromosome homology. Indeed, *PCDH11Y* has 98% nucleotide similarity to its paralogue *PCDH11X* (Supplementary Figure 3).

To date it has been very challenging to study differential protein expression between PCDH11Y and PCDH11X using immunohistochemistry because of the marked structural similarities between the proteins and limited differences in antigenicity. Studies using antibodies that do not differentiate between paralogues have been reported^26^. Others have studied gene level expression of *PCDH11Y* and *PCDH11X* in early human brain using a padlock probing and rolling circle amplification strategy^27^. Here, expression of *PCDH11Y* was found in spinal cord and medulla oblongata sections of only 46,XY human embryos (7-11 gestational weeks), and specifically in differentiating motor neuron cells. Of note, as in our data, similar levels of *PCDH11X* expression were seen in both 46,XY and 46,XX samples, whereas the expression of *PCDH11Y* appeared additive in 46,XY brain. These findings are important because they suggest that combined gene dosages of the two paralogues are *not* equal, and that *PCDH11Y* likely has a unique sex specific effect in the developing 46,XY brain.

The potential biological role of *PCDH11Y* is also relevant to sex differences. *PCDH11Y* (together with *PCDH11X*) has been implicated in cerebral hemisphere asymmetry in humans^28^. *PCDH11Y* has been associated with language development in a boy with a Y chromosome deletion involving this gene/locus^29^. A further study reported a child with severe language impairment and autistic behavior with a partial deletion of Y chromosome and consequent loss of *PCDH11Y*^30^. An additional link between *PCDH11Y* and autistic spectrum disorder (ASD) has been proposed in a small exome-wide population study^31^. Interestingly, *PCDH11Y* is unique to the human genome having evolved from the X homologue approximately 6 million years ago and it is not present in higher primates, such as chimpanzee or gorilla^32^ (Supplementary Figure 3). This finding, together with its likely role in neuronal cell communication, makes *PCDH11Y*/PCDH11Y a potentially important candidate factor to study further in relation to sex differences in human language development as well as ASD.

The other brain-specific Y chromosome gene identified was *RP11-424G14.1*. This gene encodes a long non-coding RNA (lncRNA) and is located on the long arm of the Y chromosome. Long non-coding RNA transcripts sometimes have a regulatory role on genes in the region. The gene with closest proximity to *RP11-424G14.1* is *KDM5D*, but both have a minus strand orientation and *RP11-424G14.1* is 3’ to *KDM5D*. Very little is known about any potential role of *RP11-424G14.1*, although long non-coding RNAs in general are emerging as potential regulators of transcriptomics in the human brain, and are one mechanism that might contribute to higher function in humans compared to other species^33–36^.

In addition to possible effects of sex chromosome genes, the other major mechanism of potential sex differences is through the effects of sex-dependent gonadal hormones, such as androgens (e.g. testosterone, dihydrotestosterone) or estrogens. As outlined above, the testis forms and starts to synthesize and release testosterone from around 8wpc in humans^12^. Testosterone has a direct effect on some developing tissues (e.g. Wolffian structures), whereas it is converted to the more potent hormone, dihydrotestosterone (DHT) by the enzyme 5α-reductase type 2 (encoded by *SRD5A2*), in order to have an effect on other tissues (e.g. genital tubercle). All estrogens are synthesized from testosterone by the enzyme aromatase (*CYP19A1*). The developing human ovary is believed to be relatively quiescent during early development, with limited - if any - estrogen release.

The role of sex steroids in early human brain development is still unclear. Sex steroids likely mediate several biological differences in brain development, especially in relation to sexually dimorphic regulation of endocrine systems as well as gender. The main sex hormone receptors (androgen receptor, AR; estrogen receptors, ESR1 (ERα), ESR2 (ERβ)) are more highly expressed in the hypothalamus and pituitary in adults, as well as the hippocampus^37^, and likely mediate their endocrine effects through these regions. Gender is also likely to be mediated through sex hormone receptors and androgen action, as individuals with complete androgen insensitivity syndrome (CAIS) due to impaired AR function usually have a female gender, and gender in individuals with 5α-reductase deficiency can vary^38–40^. In rodents, estrogen action is thought to play a role in sex behavior^41^, but data in humans are less convincing. The timing when all of these mechanisms are established is also unclear, although it has been suggested that sex differences occur from the mid-second trimester onwards.

Despite the lack of direct studies on human fetal brain development, *in vitro* data have recently highlighted the central role that androgen action could have on early mechanisms of brain development. Indeed, exposing human cerebral organoids to androgens (testosterone and DHT), caused an increase in cortical progenitor proliferation (that develop into excitatory cortical neurons) and an increase in the neurogenic pool^42^. Treating mouse embryonic neural stem cells with testosterone lead to sex-specific changes in gene expression; in particular an enrichment in epigenetic regulators was observed^43^. This work demonstrates how early hormonal influence potentially impacts brain development and might have long-term effects on sex-specific conditions.

Currently, very few data are available for AR expression during early human brain development, either at a transcript or protein level. In our study, we found relatively high and comparable expression of the androgen receptor gene (*AR*) in the brain cortex in both 46,XY and 46,XX fetuses at the time when the testis starts to release testosterone. These findings were supported by IHC, which showed regions of nuclear AR expression in the developing brain cortex. At a transcriptomic level, we observed a subsequent gradual decline in *AR* RNA expression in both sexes, and in both Brain-Seq 1 and Brain-Seq 2 datasets, up until the end of the observation period at 15/17wpc. Importantly, we did not detect a sex dimorphic divergence in global cortex gene expression following the onset of testicular testosterone secretion.

By IHC we were also able to find localized regions of AR expression, such as in the inferior hypothalamus. We linked this to emerging data from single cell RNA-sequencing, which identified AR expression in more localized structures such as the corticotropes of the pituitary gland^24^ (involved in adrenal and stress responses) and arcuate nucleus of the hypothalamus^25^ (implicated in appetite regulation and reproduction), although it is unclear whether sex differences occur. Other datasets, such as the Allen Brain database (https://www.brainspan.org/rnaseq/search/index.html), show some potential AR expression in the hippocampus, but transcript levels are low, and AR expression in most brain regions is low in the adult, without major sex differences (Supplementary Figure 7). Taken together, a general reduction in global AR expression in the brain cortex occurs in both sexes during the late first and early second trimesters, but subsequent localization of AR expression (and action) to key regions is likely to be important and requires further investigation.

Relatively little is known about early human fetal expression of pathways involved in conversion of testosterone to DHT (by 5α-reductase enzymes, *SRD5A*) or to estrogens (by aromatase, *CYP19A1*), or in neurosteroidogenesis (Supplementary Figures 11 and 12). Indeed, questions remain about the fetal blood brain barrier and the dynamics of secreted gonadal sex steroids reaching target tissues of interest. Again, using a global approach of brain cortex bulk RNA-Seq we were unable to detect significant expression of the key enzyme responsible for conversion of testosterone to DHT, 5α-reductase type 2 (*SRD5A2*), nor significant expression of genes involved in estrogen synthesis and action, *CYP19A1*, *ESR1* and *ESR2*. Relatively high expression of the enzymes 5α-reductase type 1 (*SRD5A1*) and 5α-reductase type 3 (*SRD5A3*) were observed in both sexes, although these have much lower affinity for the conversion of testosterone to DHT. Although several genes involved in classic and “backdoor” pathways of androgen biosynthesis were expressed, the potential “gatekeeper” in both these systems – *CYP17A1* – was not expressed. As with the androgen receptor, more localized effects in key regions or nuclei of the brain may be important.

This study has several limitations. Firstly, the anatomy of the developing cortex is complex and changes over time, and standardized sampling of fetal material is challenging. By using a bulk RNA-seq approach we have likely reduced sampling bias compared to single cell approaches, at the expense of detail, but we did generate remarkably consistent results from two independent datasets. Second, even with four samples in each karyotype group, we may have been underpowered to detect more subtle changes, although having four stages across time and in duplicated datasets should have identified major, persistent transcriptomic changes. Third, immunohistochemical analysis of *PCDH11Y* is very difficult given its similarity to *PCDH11X*. Finally, we focused mostly on global brain cortex differences over time, but more focused analysis of key regions and nuclei will provide more localized information in the future, once more specific markers are known.

Despite these limitations, this is one of the first studies to specifically address sex differences in early human fetal brain development between the late first and early second trimesters, and at a time of potentially important changes in sex hormone exposure. This work should help to focus future efforts in the field as new technologies and strategies emerge.

## Methods

### Samples and approvals

Human embryonic and fetal cerebral cortex samples used for bulk RNA-sequencing (RNA-seq) were obtained with ethical approval and with informed consent from the Medical Research Council (MRC)/Wellcome Trust-funded Human Developmental Biology Resource (HDBR) (http://www.hdbr.org) (Research Ethics Committee references: 08/H0712/34+5, 18/LO/0822, 08/H0906/21+5, 18/NE/0290; HDBR project references 200332, 200655). The HDBR is a biobank that is regulated by the UK Human Tissue Authority (HTA; www.hta.gov.uk) and acts within the HTA governance framework. The age of embryos up to 8wpc was calculated based on Carnegie staging (CS), whereas in fetuses (>8 wpc) the age was estimated from knee-heel length and foot length in relation to standard growth data. Samples were karyotyped by G-banding or quantitative PCR (chromosomes 13, 16, 18, 21, 22 and X and Y) to determine the sex of the embryo/fetus as well as to exclude any major chromosomal rearrangements or aneuploidies. Samples were stored at −70°C until RNA extraction or fixed in 10% formalin for immunohistochemistry.

Data comparing differences between 46,XX and 46,XY tissues (liver, kidney, skin, pancreas) (bulk RNA-Seq) were generated as part of a project into the developmental effects of sex chromosomes (HDBR project reference 200581) (accession number E-MTAB-13673; https://www.ebi.ac.uk/biostudies/arrayexpress/studies/0.

### Bulk RNA-Sequencing

Bulk RNA-Seq was undertaken in two independent datasets that are described in detail below. An overview of all samples included in these studies is outlined in Table 1.

#### Brain-Seq 1 dataset

In this study, total RNA was extracted from newly acquired human embryonic/fetal brain samples (n=32) using the AllPrep DNA/RNA Mini Kit (Qiagen), according to the manufacturer’s instructions. RNA quality and concentration were assessed using a Tapestation 4200 platform (Agilent, California, USA). cDNA libraries were prepared using the KAPA mRNA HyperPrep Kit (Roche) and subsequently sequenced on a NextSeq 500 sequencer (paired-end 43 base pairs) (Illumina, San Diego, CA) in a single run to reduce potential batch effects. Fastq files were processed by FastQC (Babraham Bioinformatics) and aligned to the Human Genome (Ensembl, GRCh37) using STAR (2.5.2a)^44^. The matrix containing uniquely mapped read counts was generated using featureCounts^45^, part of the R package Rsubread. Differential-expression analysis was performed using DESeq2^46^. A cut-off adjusted p-value of 0.05 and log2 fold change of 0.7 were used.

#### Brain-Seq 2 dataset

Bulk RNA-seq data for an independent parallel replication study (Brain-Seq 2) were obtained from a Human Developmental Biology Resource (HDBR) project focusing on early human brain development (https://www.ebi.ac.uk/gxa/experiments/E-MTAB-4840/Downloads)^17^, Data were analyzed using the bioinformatic pipeline described above. To achieve a balanced experimental design and match the developmental stages to those used in the Brain-Seq 1 dataset, n=8 samples per stage (four 46,XX, four 46,XY) were included from the Brain-Seq 2 dataset (Table 1).

### Quantitative reverse transcriptase PCR (qRT-PCR)

Purified RNA was quantified using a NanoDrop 1000 spectrophotometer (Thermo Fisher Scientific). RNA (2 μg) was reverse transcribed with a SuperScript III Reverse Transcription kit (Applied Biosystems, Thermo Fisher Scientific) according to the manufacturer’s instructions. qRT-PCR was performed using TaqMan Fast Advanced Master Mix (Applied Biosystems) and TaqMan gene assays (*AR* Hs00171172_m1; *PCDH11Y* Hs00263145_m1, Thermo Fisher Scientific) on a StepOnePlus System (Thermo Fisher Scientific). The relative expression of gene was calculated as 2–ΔΔCt using the comparative Ct (ΔΔCt) method and GAPDH (Hs02758991_g1) (Thermo Fisher Scientific) as an internal housekeeping control. Experiments were repeated on two independent occasions; three samples per developmental stage were used, each performed with triplicate replicates. Four control tissues were used (kidney, liver, pancreas, skin). Representative data of one experiment are shown.

### Adult gene expression data

Expression data for key genes of interest in adult tissues was obtained from the Human Protein Atlas (HPA) Consensus expression data (v23) (https://www.proteinatlas.org/)^5^ and from the Genotype-Tissue Expression (GTEx) Project (https://gtexportal.org/home/). The data used for the analyses described in this manuscript were obtained from the HPA and GTEx Portal on 10/10/23.

### Genomic characterization

The genomic location and structure of genes of interest was obtained from Ensembl Genome Browser 110 accessed on 10/10/23 (https://www.ensembl.org/index.html) (GRCh38.p14)^47^. Species comparisons for PCDH11Y were generated using GeneTree (ENSGT00940000158335) in the Ensembl Genome Browser.

### Immunohistochemistry (IHC)

Human brain cortex samples (46,XY) at 10wpc and 17wpc were fixed in 4% paraformaldehyde before being processed, embedded and sectioned for histology and immunohistochemistry (IHC). Hematoxylin and eosin (H&E) staining was performed using standard methods on 3µm sections. IHC was performed on a Leica Bond-max automated platform (Leica Biosystems). In brief, antigen retrieval was undertaken to unmask the epitope (Heat Induced Epitope Retrieval (HIER), Bond-max protocol F), then endogenous activity was blocked with peroxidase using a Bond polymer refine kit (cat # DS9800). Next, slides were incubated with a primary androgen receptor (AR, NR3C4) antibody for 1 hour (Abcam ab108341 ChIP grade, 1:50 dilution, HIER2 for 20 mins) before having a post-primary antibody applied and horseradish peroxidase (HRP) labelled polymer, followed by 3, 3-diaminobenzidine (DAB) chromogen solution (all Bond polymer refine kit). Sections were counterstained with hematoxylin, washed, dehydrated in graded alcohols, cleared in two xylene changes and mounted. Imaging was undertaken using an Aperio CS2 Scanner (Leica Biosystems) at 40x objective. Subsequent analysis was performed with QuPath (v.0.2.3) (https://qupath.github.io) and Leica ImageScope (Leica Biosystems) software.

### Single cell RNA-sequencing (scRNA-Seq) data

scRNA-Seq data were obtained for androgen receptor (*AR*) expression using the following resources:

1. human fetal pituitary cell images were generated from the open resource (CC BY 4.0 license) on the shiny webpage https://tanglab.shinyapps.io/Human_Fetal_Pituitary_Endocrine_Cells/ developed by Zhang et al.^24^;
2. human fetal hypothalamus images were generated from the NeMo Analytic open resource (CC-BY-NC 4.0 International license) webpage https://nemoanalytics.org/p?l=a856c14e&g=gad2 developed by Herb et, al.^25^.

Data are represented as UMAP (uniform manifold approximation and projection, for dimension reduction) images for cell clusters at different stages of development.

### Graphical representation

Venn diagrams were produced using the online tool InteractiVenn (http://www.interactivenn.net/#) ^48^. Graphics were generated using GraphPad Prism version 8.4.3 for Windows (GraphPad Software, San Diego, USA; www.graphpad.com).

### Statistical analysis

Statistical analyses were performed using GraphPad Prism version 8.4.3 for Windows (GraphPad Software, San Diego, USA; www.graphpad.com) and R (RStudio 2022.07.2 for macOS), and are described in the relevant figure legends. A p-value <0.05 was considered significant.

## Data Availability

The datasets generated and analyzed during the current study can be found in the following links:

1. Brain-Seq1 (bulk RNA-seq): ArrayExpress/Biostudies (accession number E-MTAB-13662; https://www.ebi.ac.uk/biostudies/arrayexpress/studies)
2. Brain-Seq2 (bulk RNA-seq): https://www.ebi.ac.uk/gxa/experiments/E-MTAB-4840/Downloads^17^
3. Control samples (bulk RNA-seq): ArrayExpress/Biostudies (accession number E-MTAB-13673; https://www.ebi.ac.uk/biostudies/arrayexpress/studies)

## Acknowledgements

This research was funded in whole, or in part, by the Wellcome Trust (grant 209328/Z/17/Z). For the purpose of Open Access, the author has applied a CC BY public copyright license to any Author Accepted Manuscript version arising from this submission. We also thank other members of the Human Developmental Biology Resource and UCL Genomics for their additional contributions to this work. Human fetal material was provided by the Joint MRC/Wellcome Trust (Grant MR/R006237/1, MR/X008304/1 and 226202/Z/22/Z) Human Developmental Biology Resource (http://www.hdbr.org).

Research at UCL GOS Institute of Child Health has support from the National Institute for Health Research, Great Ormond Street Hospital Biomedical Research Centre (grant IS-BRC-1215-20012). The views expressed are those of the authors and not necessarily those of the National Health Service, National Institute for Health Research, or Department of Health.

The Genotype-Tissue Expression (GTEx) Project was supported by the Common Fund of the Office of the Director of the National Institutes of Health, and by NCI, NHGRI, NHLBI, NIDA, NIMH, and NINDS. The data used for the analyses described in this manuscript were obtained from the GTEx Portal on 10/10/23.

## Author Contributions

Author contributions were as follows. Study conceptualization: FB, IdV, JCA; Methodology: FB, JPS, IdV, JCA; Investigation: FB, JPS, OKO, AJ, NM, PN, TB, NS, MTD, IdV; Formal analysis: FB, JPS, IdV; Data curation: FB, IdV; Resources: NS; Project administration: FB, JCA; Supervision: FB, IdV, JCA; Validation: FB, IdV, JCA; Visualization: FB, OKO, JCA; Writing – original draft: FB, JCA; Writing – review & editing: All authors; Funding acquisition: JCA.

## Supplementary information

### Supplementary Table 1

### Supplementary Figures 1-15

### Supplementary Data1_Brain-Seq 1

- Supplementary data 1.1. Samples used in Brain-Seq 1 dataset.
- Supplementary data 1.2. Top 250 differentially expressed genes, 46,XY versus 46,XX at CS22-23.
- Supplementary data 1.3. Top 250 differentially expressed genes, 46,XX versus 46,XY at CS22-23.
- Supplementary data 1.4. Top 250 differentially expressed genes, 46,XY versus 46,XX at 9wpc.
- Supplementary data 1.5. Top 250 differentially expressed genes, 46,XX versus 46,XY at 9wpc.
- Supplementary data 1.6. Top 250 differentially expressed genes, 46,XY versus 46,XX at 11-12wpc.
- Supplementary data 1.7. Top 250 differentially expressed genes, 46,XX versus 46,XY at 11-12wpc.
- Supplementary data 1.8. Top 250 differentially expressed genes, 46,XY versus 46,XX at 15-17wpc.
- Supplementary data 1.9. Top 250 differentially expressed genes, 46,XX versus 46,XY at 15-17wpc.
- Supplementary data 1.10. Differentially expressed genes, 46,XX 9wpc versus CS22-23.
- Supplementary data 1.11. Differentially expressed genes, 46,XY 9wpc versus CS22-23.
- Supplementary data 1.12. Differentially expressed genes, 46,XX 15-17wpc versus CS22-23.
- Supplementary data 1.13. Differentially expressed genes, 46,XY 15-17wpc versus CS22-23.

#### Supplementary Data2_Brain-Seq 2

- Supplementary data 2.1. Samples used in Brain-Seq 2 dataset.
- Supplementary data 2.2. Top 250 differentially expressed genes, 46,XY versus 46,XX at CS22-23.
- Supplementary data 2.3. Top 250 differentially expressed genes, 46,XX versus 46,XY at CS22-23.
- Supplementary data 2.4. Differentially expressed genes, 46,XY versus 46,XX at 9wpc.
- Supplementary data 2.5. Top 250 differentially expressed genes, 46,XX versus 46,XY at 9wpc.
- Supplementary data 2.6. Top 250 differentially expressed genes, 46,XY versus 46,XX at 11-12wpc.
- Supplementary data 2.7. Top 250 differentially expressed genes, 46,XX versus 46,XY at 11-12wpc.
- Supplementary data 2.8. Differentially expressed genes, 46,XY versus 46,XX at 15-17wpc.
- Supplementary data 2.9. Differentially expressed genes, 46,XX versus 46,XY at 15-17wpc.
- Supplementary data 2.10. Differentially expressed genes, 46,XX 9wpc versus CS22-23.
- Supplementary data 2.11. Differentially expressed genes, 46,XY 9wpc versus CS22-23.
- Supplementary data 2.12. Differentially expressed genes, 46,XX 15-17wpc versus CS22-23.
- Supplementary data 2.13. Differentially expressed genes, 46,XY 15-17wpc versus CS22-23.

#### Supplementary Data3_Controls

- Supplementary data 3.1. Control samples used.
- Supplementary data 3.2. Top 250 differentially expressed genes, skin samples 46,XY versus 46,XX at CS22-23.
- Supplementary data 3.3. Differentially expressed genes, skin samples 46,XX versus 46,XY at CS22-23.
- Supplementary data 3.4. Differentially expressed genes, pancreas samples 46,XY versus 46,XX at 9wpc.
- Supplementary data 3.5. Differentially expressed genes, pancreas samples 46,XX versus 46,XY at 9wpc.
- Supplementary data 3.6. Differentially expressed genes, liver samples 46,XY versus 46,XX at 11-12wpc.
- Supplementary data 3.7. Differentially expressed genes, liver samples 46,XX versus 46,XY at 11-12wpc.
- Supplementary data 3.8. Differentially expressed genes, kidney samples 46,XY versus 46,XX at 15-17wpc.
- Supplementary data 3.9. Differentially expressed genes, kidney samples 46,XX versus 46,XY at 15-17wpc.

