## Supplementary Material for "Sex differences in early human fetal brain development"

### Supplementary Table 1

| **Brain-enriched Y chromosome genes** | **RNA tissue specificity (consensus dataset)** |
|---|---|
| *AC009977.1* | na |
| *AC010970.2* | na |
| *ACTG1P2* | na |
| *AGKP1* | na |
| *AMELY* | not detected |
| *ANKRD36P1* | na |
| *ANOS2P* | na |
| *ARSFP1* | na |
| *BCORP1* | na |
| *CD99* | Low tissue specificity |
| *DDX3Y* | Tissue enhanced (bone marrow) |
| *EIF1AY* | Tissue enhanced (heart muscle, tongue) |
| *FAM8A4P* | na |
| *GPR143P* | na |
| *GYG2P1* | na |
| *HSFY2* | Tissue enriched (testis) |
| *HSFY3P* | na |
| *KDM5D* | Low tissue specificity |
| *MED14P1* | na |
| *NAP1L1P2* | na |
| *NLGN4Y* | Tissue enhanced (seminal vesicle) |
| *P2RY8* | Group enriched (bone marrow, lymphoid tissue) |
| *PARPAP1* | na |
| *PCDH11Y* | Tissue enhanced (brain) |
| *PRKY* | na |
| *RFTN1P1* | na |
| *RNU2-57P* | na |
| *RP11-424G14.1* | na |
| *RP11-576C2.1* | na |
| *RPS4Y1* | Low tissue specificity |
| *RPS4Y2* | Group enriched (prostate, testis) |
| *SHOX* | Low tissue specificity |
| *SRY* | Tissue enhanced (skin, testis) |
| *TAB3P1* | na |
| *TBL1Y* | Tissue enhanced (prostate, thyroid gland) |
| *TMSB4Y* | Tissue enhanced (intestine) |
| *TPTE2P4* | na |
| *TSPY2* | Tissue enriched (testis) |
| *TTTY10* | na |
| *TTTY14* | na |
| *TTTY15* | na |
| *TXLNGY* | na |
| *USP9Y* | Low tissue specificity |
| *USP9YP14* | na |
| *USP9YP28* | na |
| *UTY* | Low tissue specificity |
| *XG* | Tissue enhanced (adipose tissue, skin) |
| *ZFY* | Low tissue specificity |
| *ZFY-AS1* | na |

**Supplementary Table 1. Expression of Y chromosome genes in human adult tissues.** Data were obtained from the Human Protein Atlas “Consensus” dataset for bulk RNA-sequencing (50 tissues, including 10 brain/neuronal regions). The brain specific expression of *PCDH11Y* is indicated in red. na, not available.

### Supplementary Figure 1

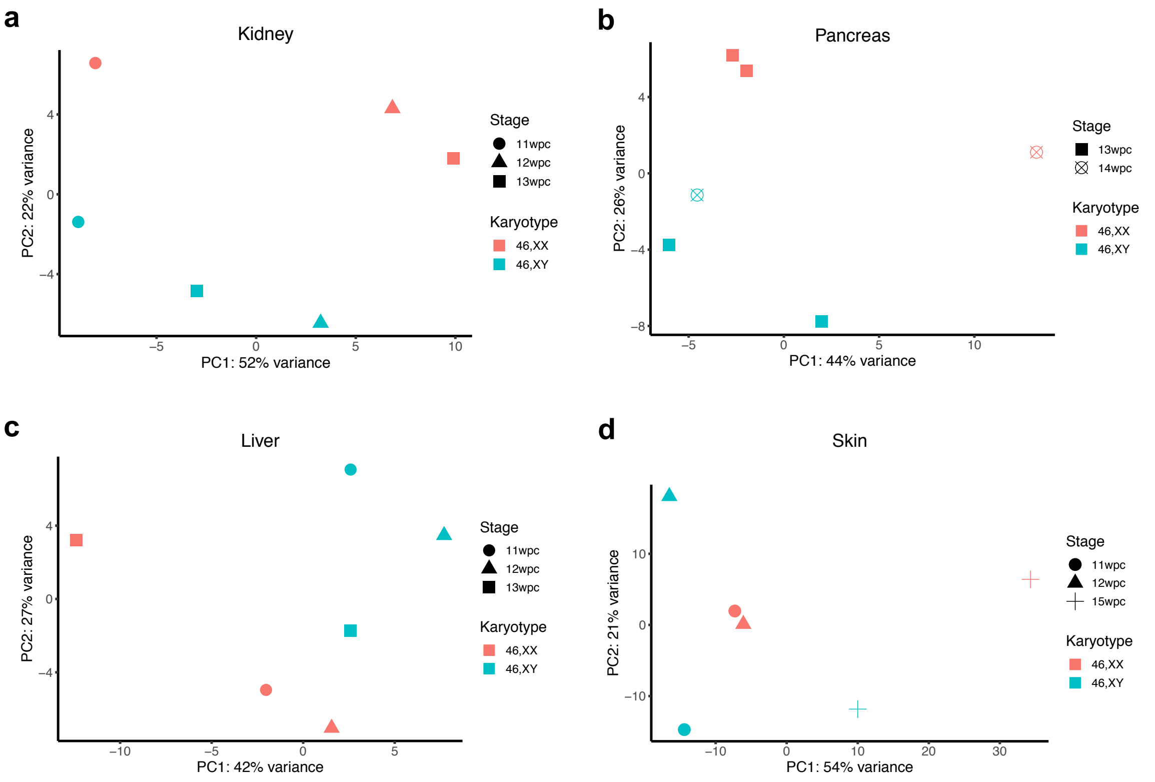


**Supplementary Figure 1. Principal component analysis (PCA) showing effect of karyotype in non-brain control tissues.** Data for bulk RNA-sequencing of kidney, pancreas, liver and skin are shown (n=6 each group). Three 46,XX and three 46,XY samples were included for each tissue, matched for developmental stage, as indicated. PC, principal component.

### Supplementary Figure 2

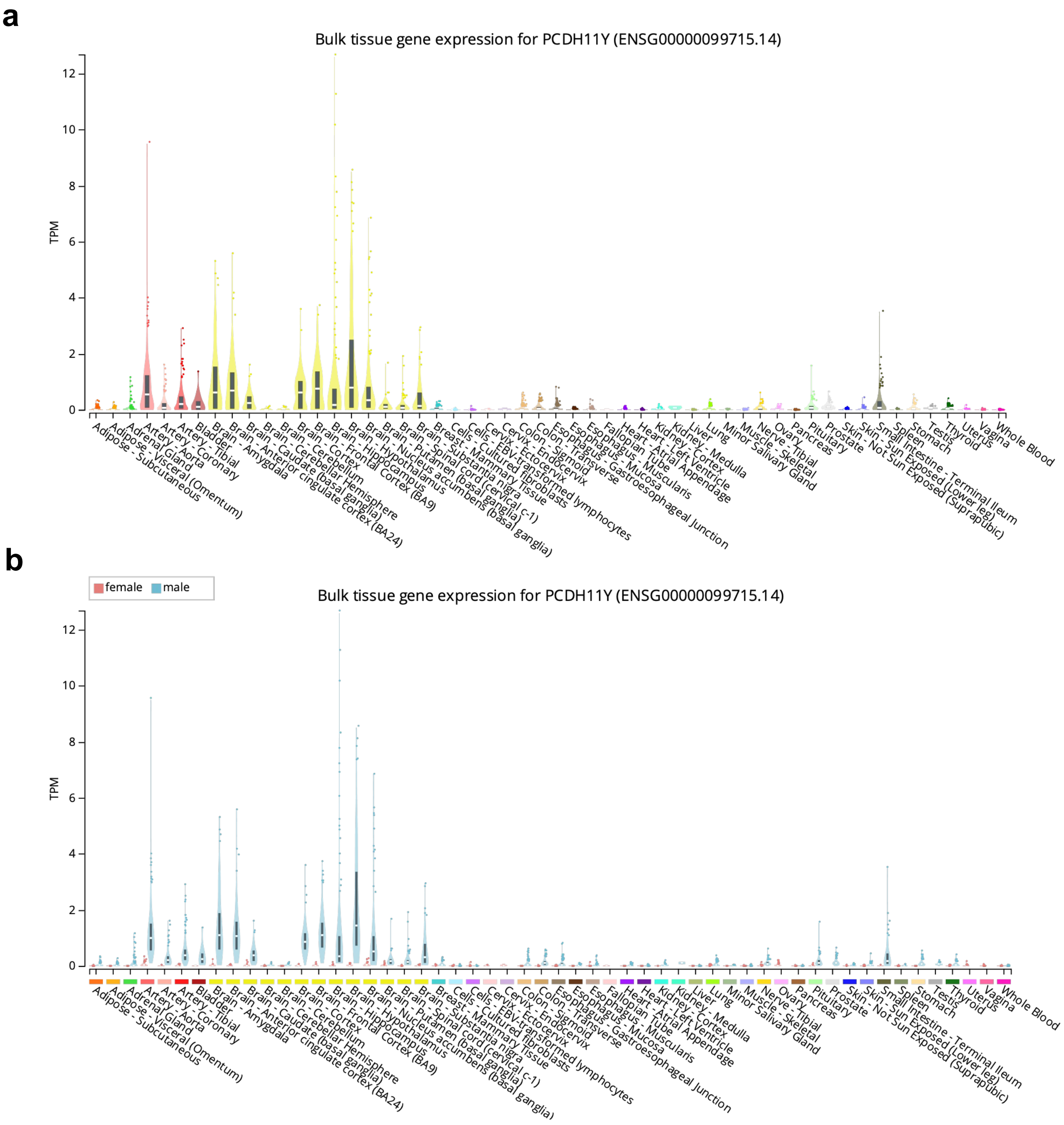


**Supplementary Figure 2. Differential expression of *PCDH11Y* in a panel of adult tissues. a** *PCDH11Y* expression in all samples. **b** *PCDH11Y* expression shown by sex. Data represent bulk RNA-sequencing from the Genotype-Tissue Expression (GTEx) Project. GTEx was supported by the Common Fund of the Office of the Director of the National Institutes of Health, and by NCI, NHGRI, NHLBI, NIDA, NIMH, and NINDS. The data were obtained from the GTEx Portal on 10/10/23 (GTEx Analysis Release V8; dbGaP Accession phs000424.v8.p2). TPM, transcripts per million.

### Supplementary Figure 3

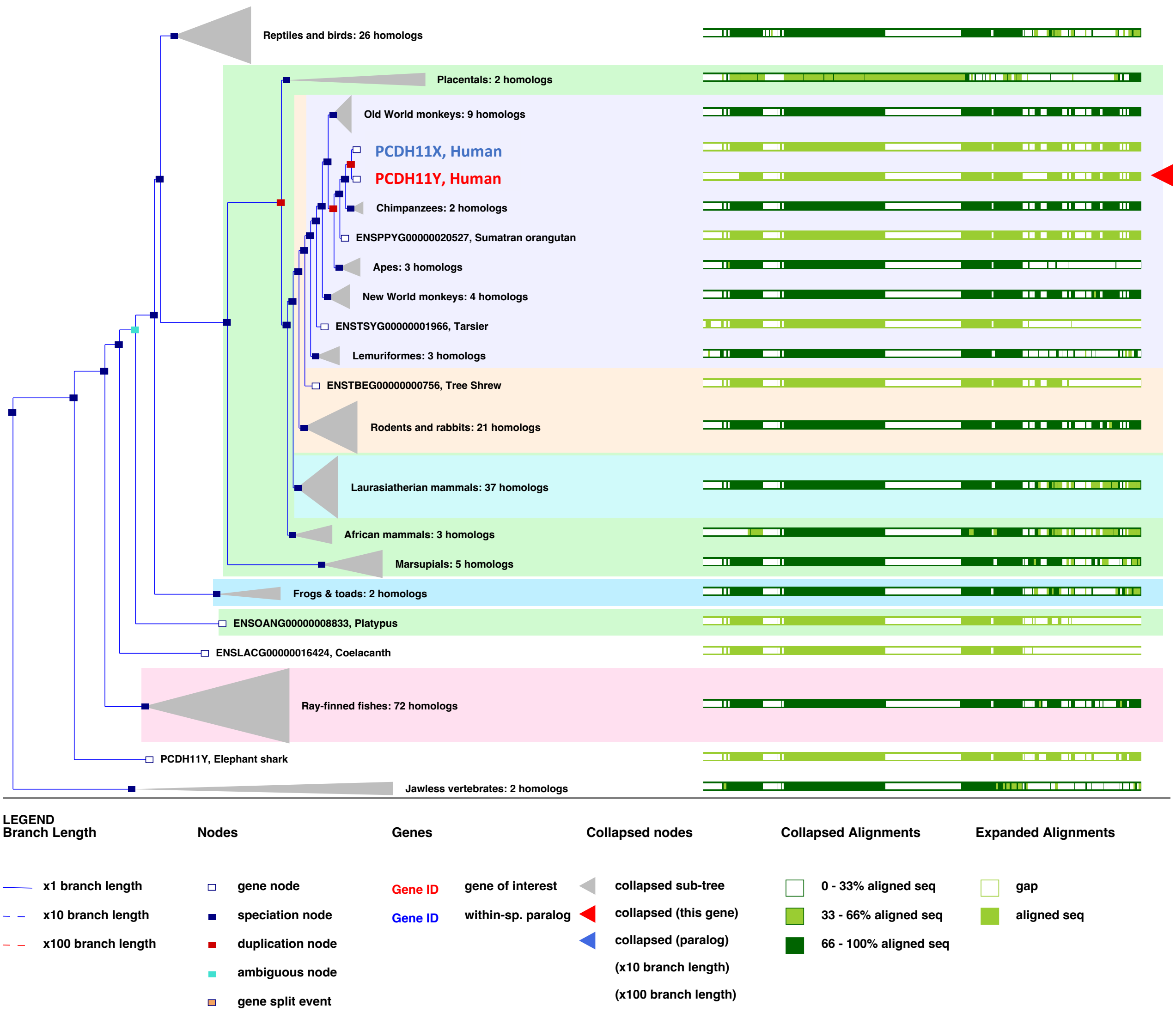


**Supplementary Figure 3. Comparative genomics of *PCDH11X*/*PCDH11Y* in humans with 185 speciation nodes.** The human *PCDH11X* transcript is shown in blue and has strong alignment with key domains of the paralogue in other species. The human *PCDH11Y* transcript is shown in red and shows localized regions of variance. Data derived and reproduced from GeneTree (ENSGT00940000158335) in Ensembl Genome Browser release 110 (https://www.ensembl.org/) (Martin FJ, et al. Nucleic Acids Res. 2023, 51(D1):D933-D941; doi:10.1093/nar/gkac958).

### Supplementary Figure 4

**a**

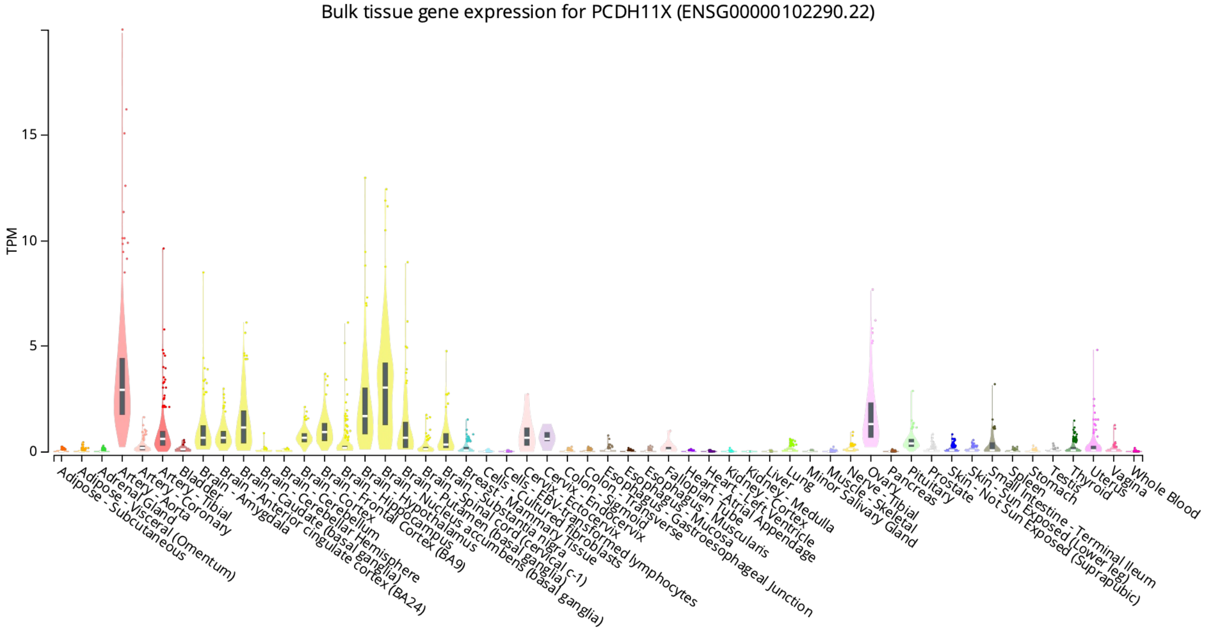


**b**

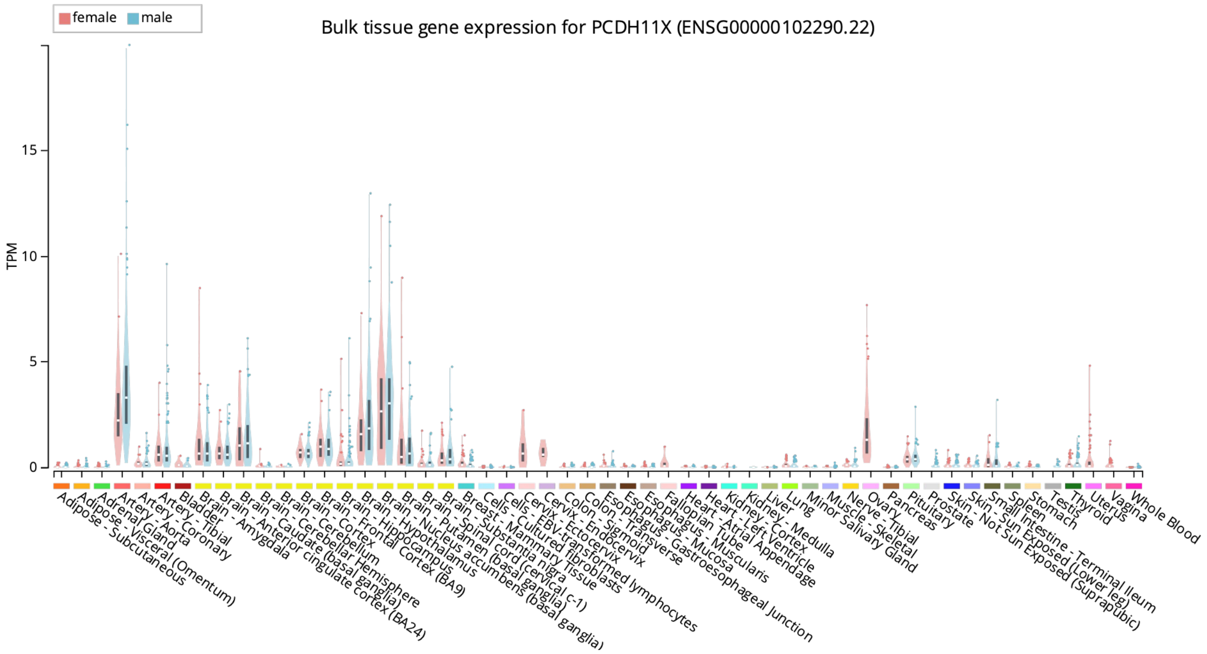


**Supplementary Figure 4. Differential expression of *PCDH11X* in a panel of adult tissues. a** *PCDH11X* expression in all samples. **b** *PCDH11X* shown by sex. Data represent bulk RNA-sequencing from the Genotype-Tissue Expression (GTEx) Project. GTEx was supported by the Common Fund of the Office of the Director of the National Institutes of Health, and by NCI, NHGRI, NHLBI, NIDA, NIMH, and NINDS. The data were obtained from the GTEx Portal on 10/10/23 (GTEx Analysis Release V8; dbGaP Accession phs000424.v8.p2). TPM, transcripts per million.

### Supplementary Figure 5

a

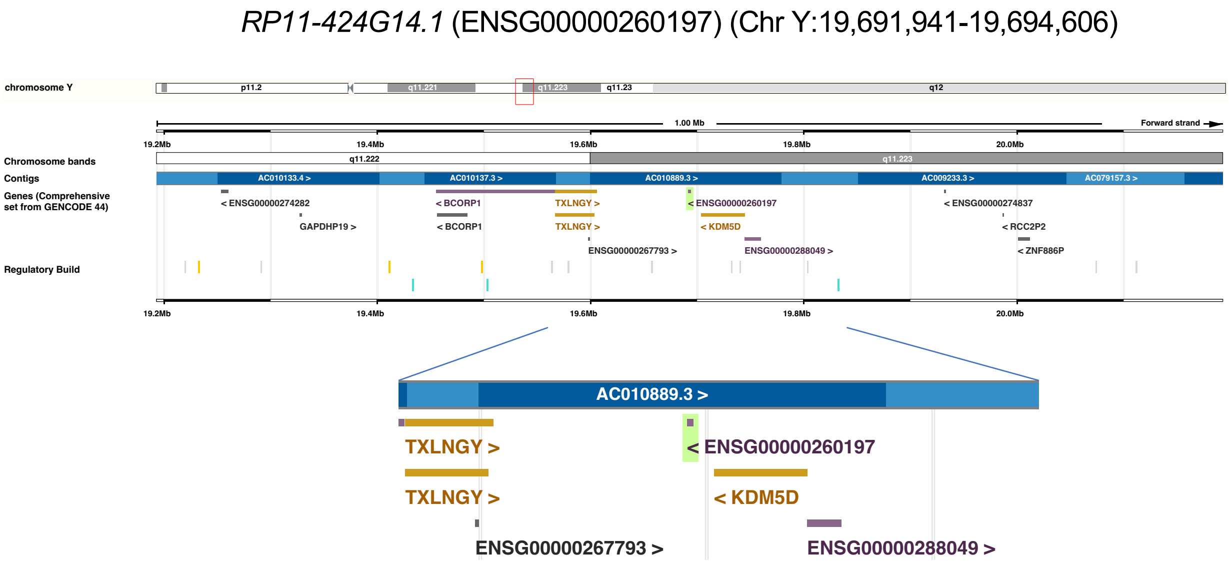


b

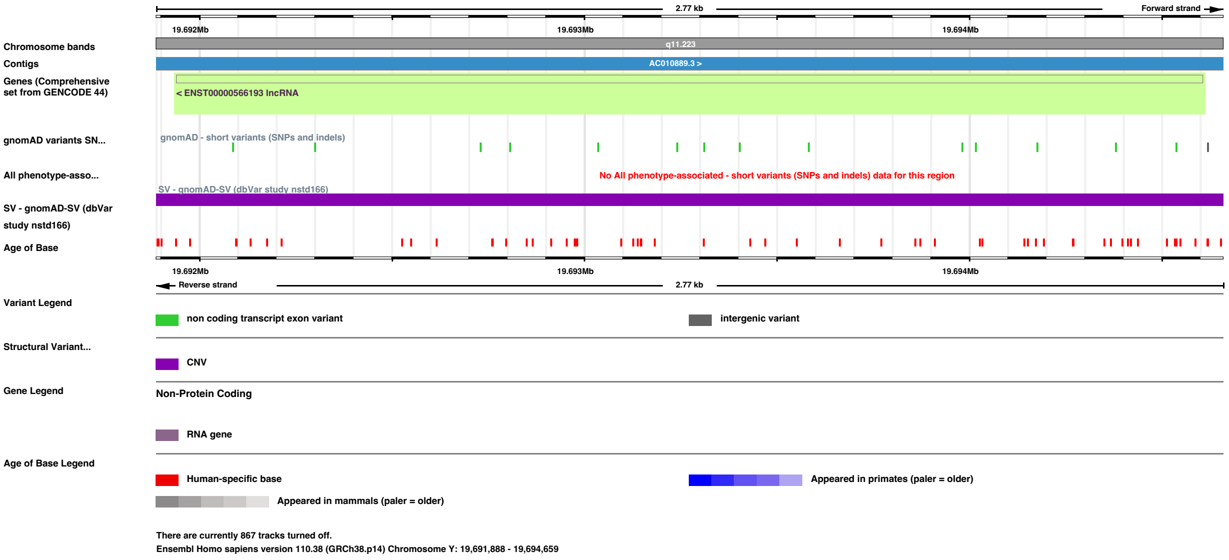


**Supplementary Figure 5. Genomic locus of the long non-coding RNA, *RP11-424G14.1*. a** Location of *RP11-424G14.1* (shown as Ensembl gene ENSG00000260197) on the long arm of the Y chromosome in minus orientation and centromeric to *KDM5D*. b Locus characteristics of the transcript (ENST00000566193.1) associated with *RP11-424G14.1.* Data derived and reproduced from Ensembl release 110 (https://www.ensembl.org/) (Martin FJ, et al. Nucleic Acids Res. 2023, 51(D1):D933-D941; doi:10.1093/nar/gkac958).

### Supplementary Figure 6

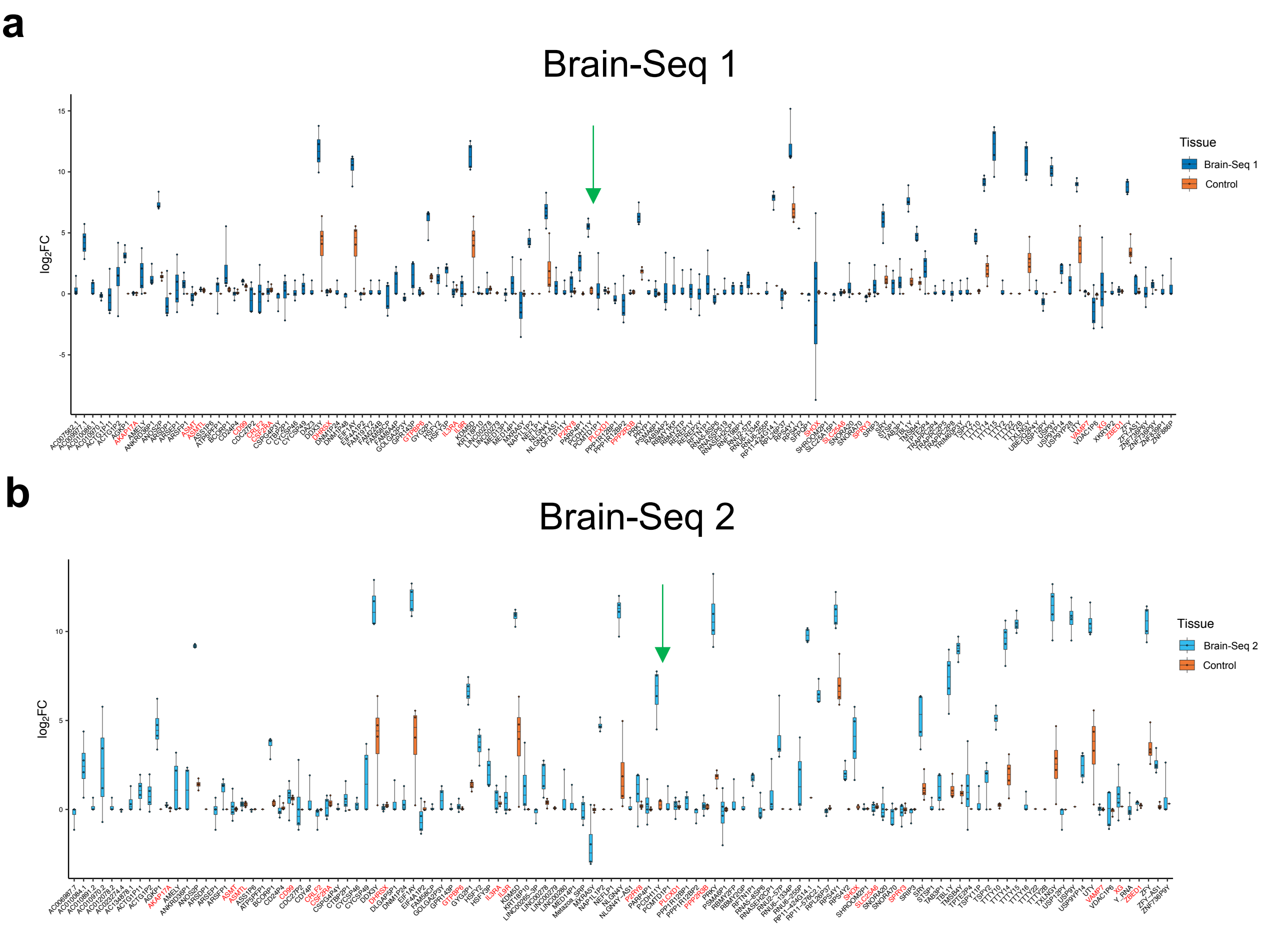


**Supplementary Figure 6. Expression of Y chromosome genes in human fetal brain cortex samples compared to control tissues. a** Brain-Seq 1 dataset. **b** Brain-Seq 2 dataset**.** Data are shown for bulk RNA-sequencing as log2 fold change (FC) of Y chromosome genes following comparison of 46,XY vs 46,XX brain samples and 46,XY vs 46,XX control samples. Each brain dataset consisted of n=32 cortex samples between CS22/23 and 15-17 weeks post conception (wpc). Control dataset consisted of n=32 samples including skin, pancreas, liver and kidney tissues between 11 and 15 wpc (each tissue having n=8). Boxplots represent the distribution of log2FC values of each gene, with the box extending from the first to the third quartile and the median shown with a line. Genes labelled in red are in pseudoautosomal regions (PAR). FC, fold change. Green arrows indicate the position of *PCDH11Y.*

### Supplementary Figure 7

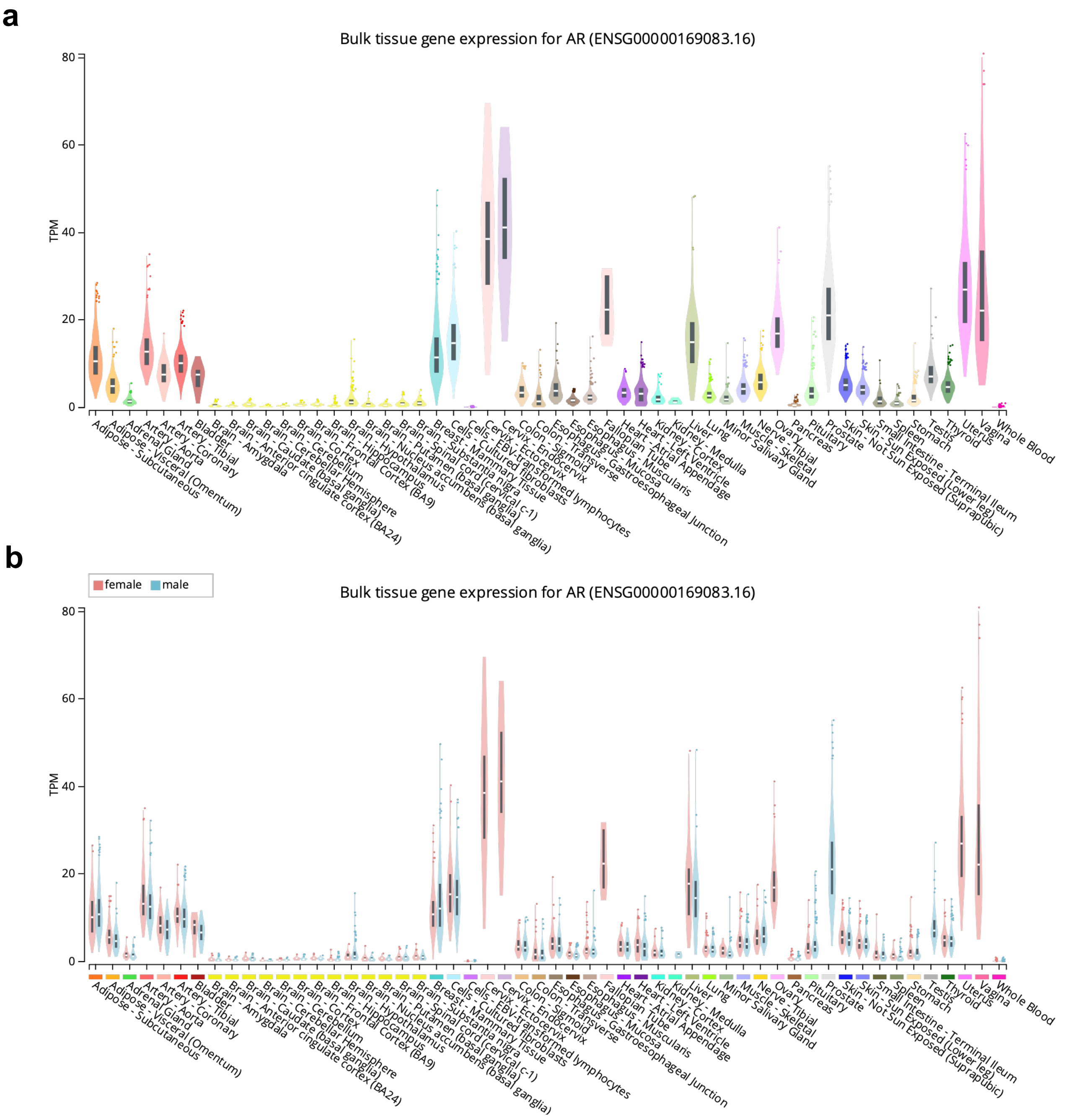


**Supplementary Figure 7. Differential expression of the gene encoding the androgen receptor (*AR*) in a panel of adult tissues. a** *AR* expression in all samples. **b** *AR* expression shown by sex. Data represent bulk RNA-sequencing from the Genotype-Tissue Expression (GTEx) Project. GTEx was supported by the Common Fund of the Office of the Director of the National Institutes of Health, and by NCI, NHGRI, NHLBI, NIDA, NIMH, and NINDS. The data were obtained from the GTEx Portal on 10/10/23 (GTEx Analysis Release V8; dbGaP Accession phs000424.v8.p2). TPM, transcripts per million.

#### Supplementary Figure 8

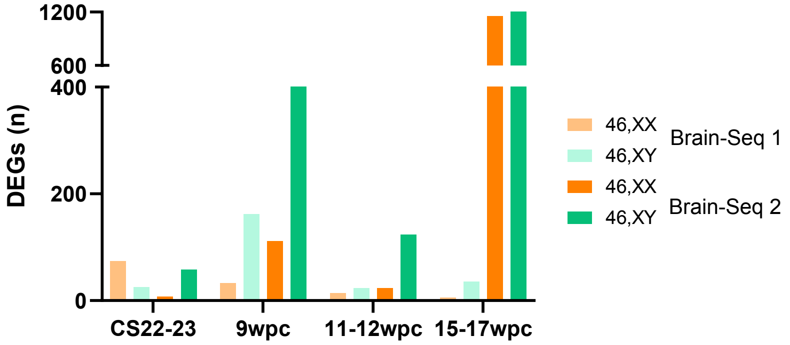


**Supplementary Figure 8. Number of differentially expressed genes (DEGs) (46,XX versus 46,XY) in the Brain-Seq 1 and Brain-Seq 2 datasets with developmental stage. a** All DEGs in both 46,XX and 46,XY samples. A cut off for differential expression of log2 fold change >0.7 and p-adj <0.05 was used. DEGs, differentially expressed genes; wpc, weeks post conception

### Supplementary Figure 9

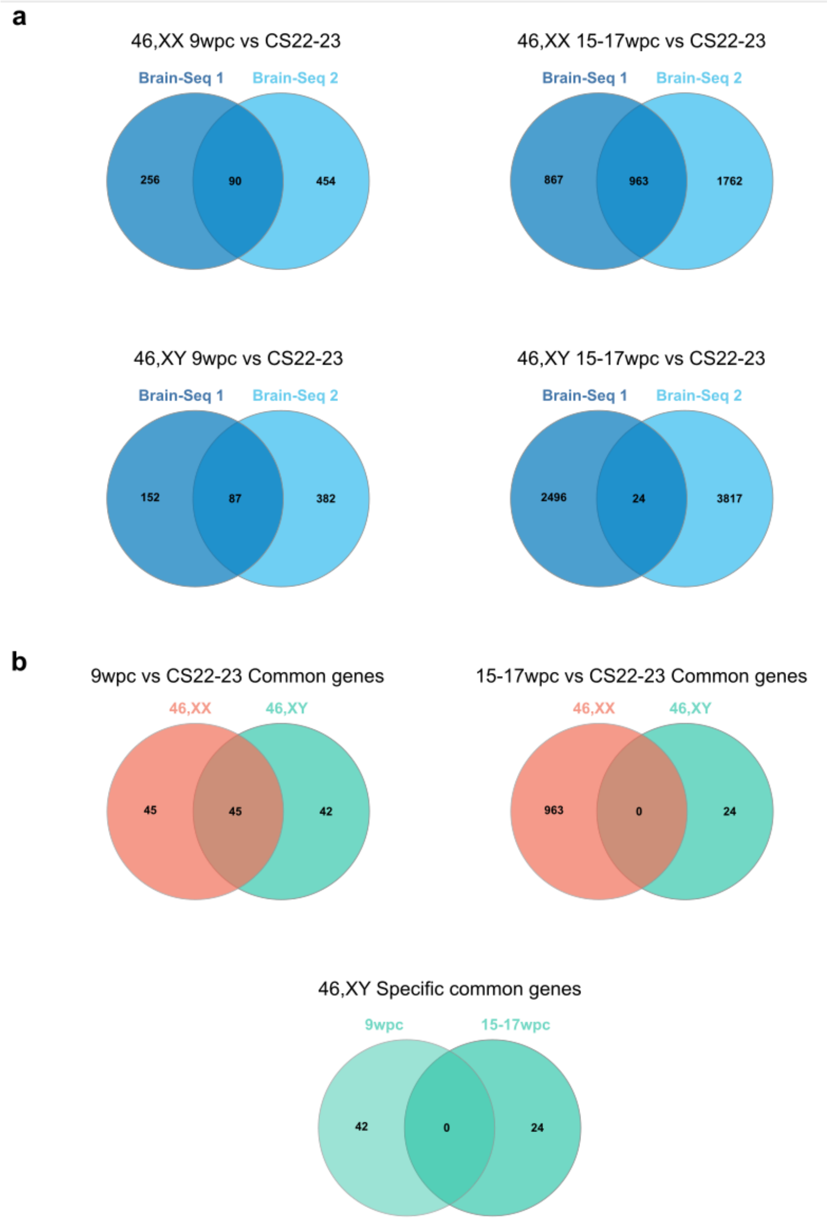


**Supplementary Figure 9. Androgen responsive time series analyses of differentially expressed genes (DEGs) in Brain-Seq 1 and Brain-Seq 2 datasets. a** DEGs in both 46,XX and 46,XY comparing 9wpc and CS22-23, and 15-17wpc and CS22-23. **b** DEGs common to both Brain-Seq datasets in 9wpc versus CS22-23 (46,XX 9wpc versus CS22-23, n=90; 46,XY 9wpc versus CS22-23, n=87) and 15-17wpc versus CS22-23 (46,XX 15-17wpc versus CS22-23, n=963; 46,XY 15-17wpc versus CS22-23, n=24). No 46,XY specific common DEGs were identified. vs, versus.

#### Supplementary Figure 10

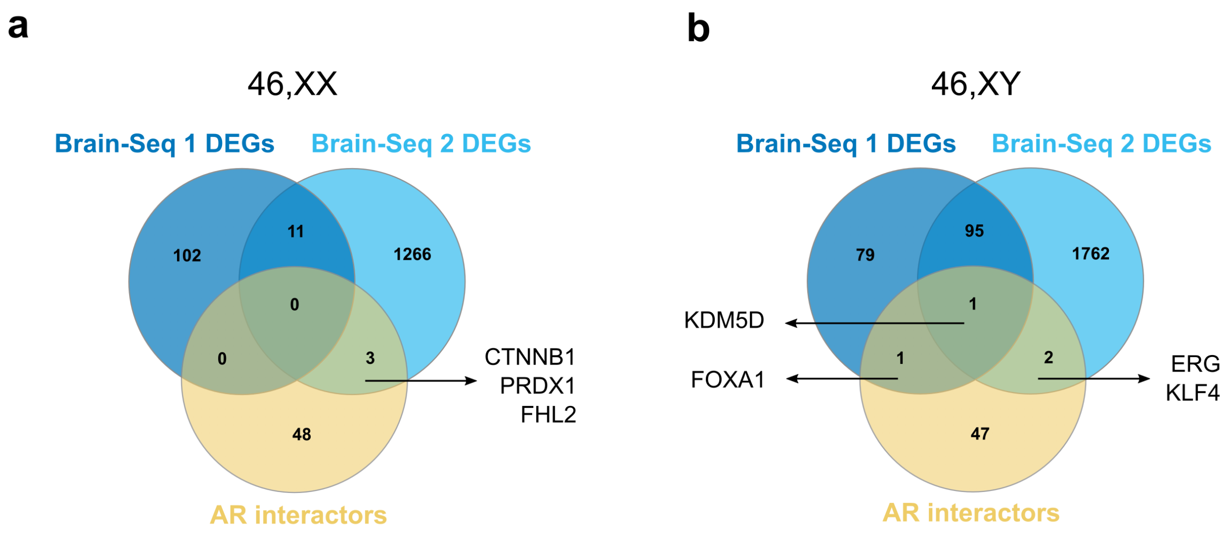


**Supplementary Figure 10. Venn diagrams showing the intersection of AR interactors with Brain-Seq differentially expressed genes (DEGs). a** Overlap of DEGs in both 46,XX datasets with AR interactors. **b** Overlap of DEGs in both 46,XX datasets with AR interactors. AR interactors are reported proteins that interact with the androgen receptor as listed in the Reactome Knowledgebase (https://reactome.org) (Gillespie et al., 2022).

### Supplementary Figure 11

a

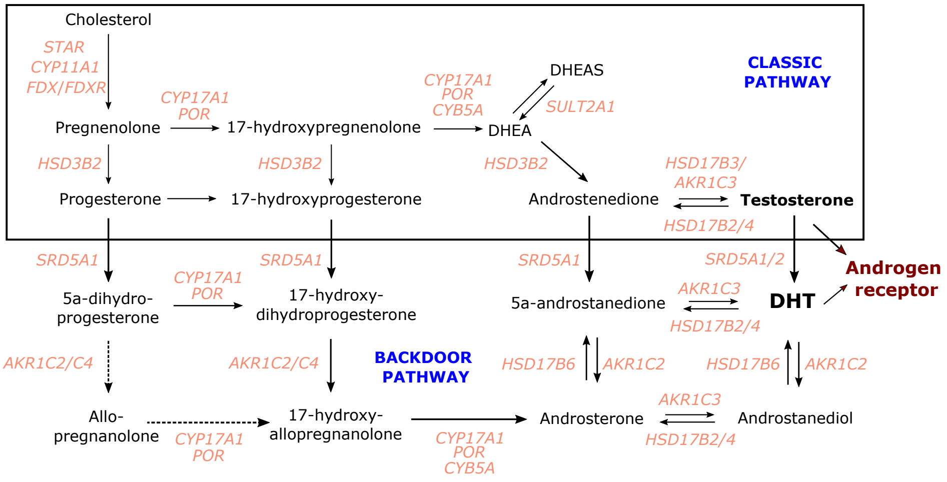


b

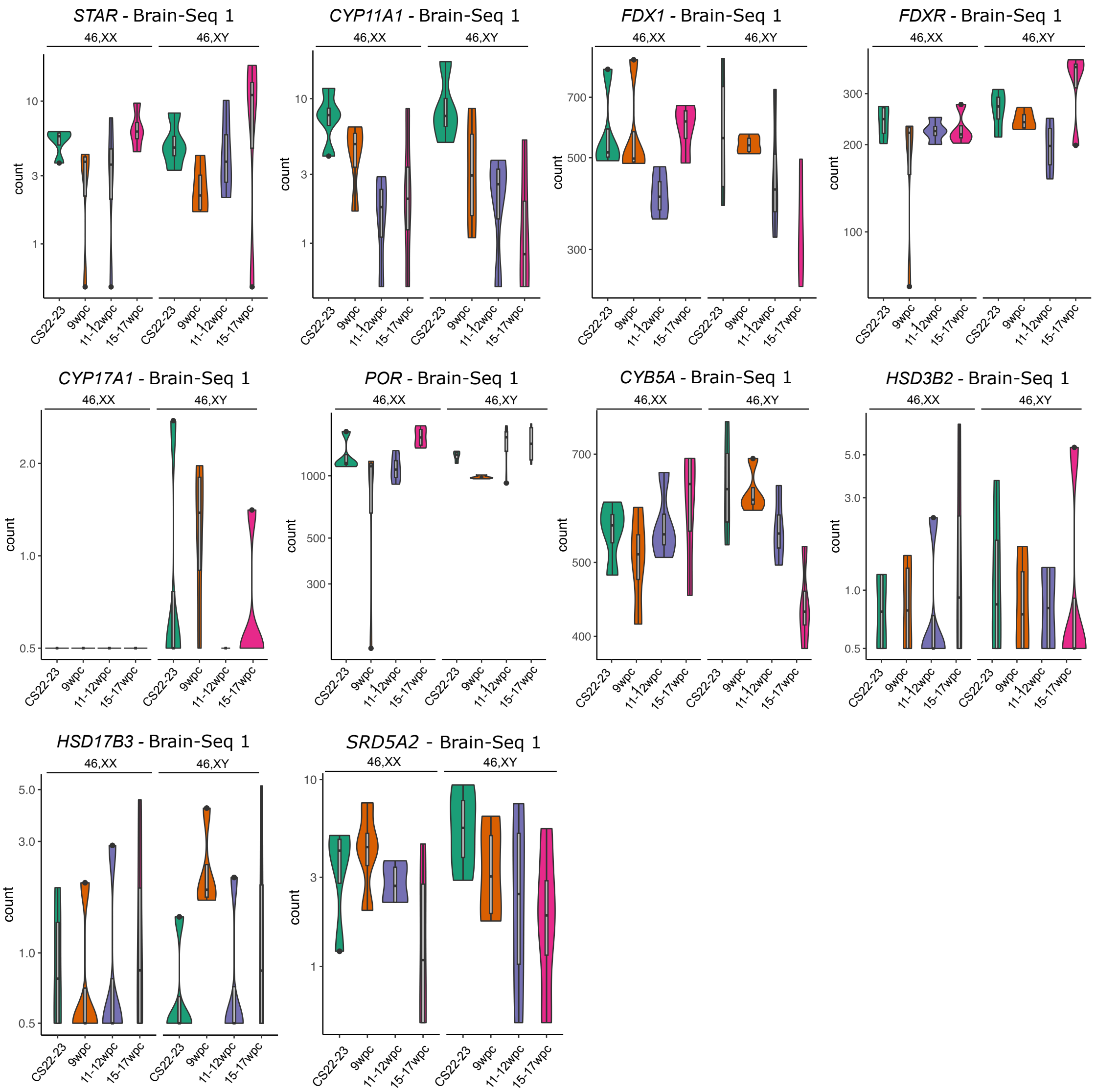

**Supplementary Figure 11. Expression of genes involved in the classic steroid pathway in the human brain cortex during early fetal development. a** Classic steroid biosynthesis pathway. Adapted from del Valle I et al., JCI Insight 2023;8(14):e168177; doi.org/10.1172/jci.insight.168177. **b** Data are shown as violin plots of normalized counts on a log scale, with n=4 in each group. CS, Carnegie stage; wpc, weeks post-conception.

### Supplementary Figure 12

**a**

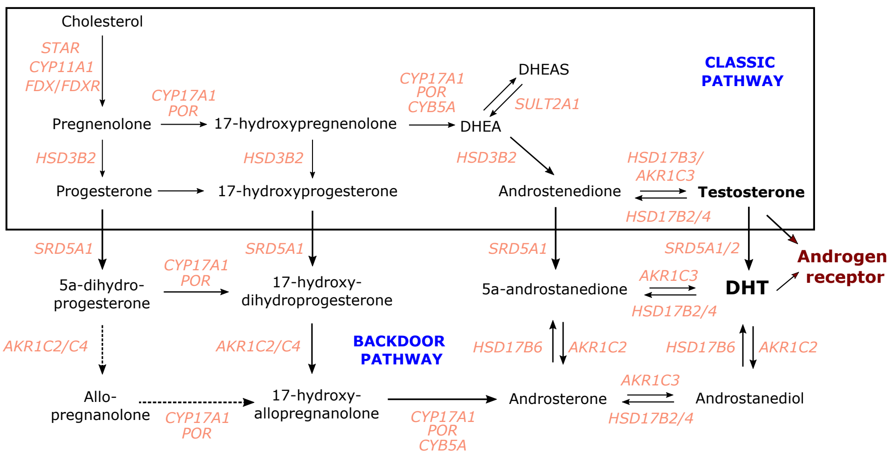


**b**

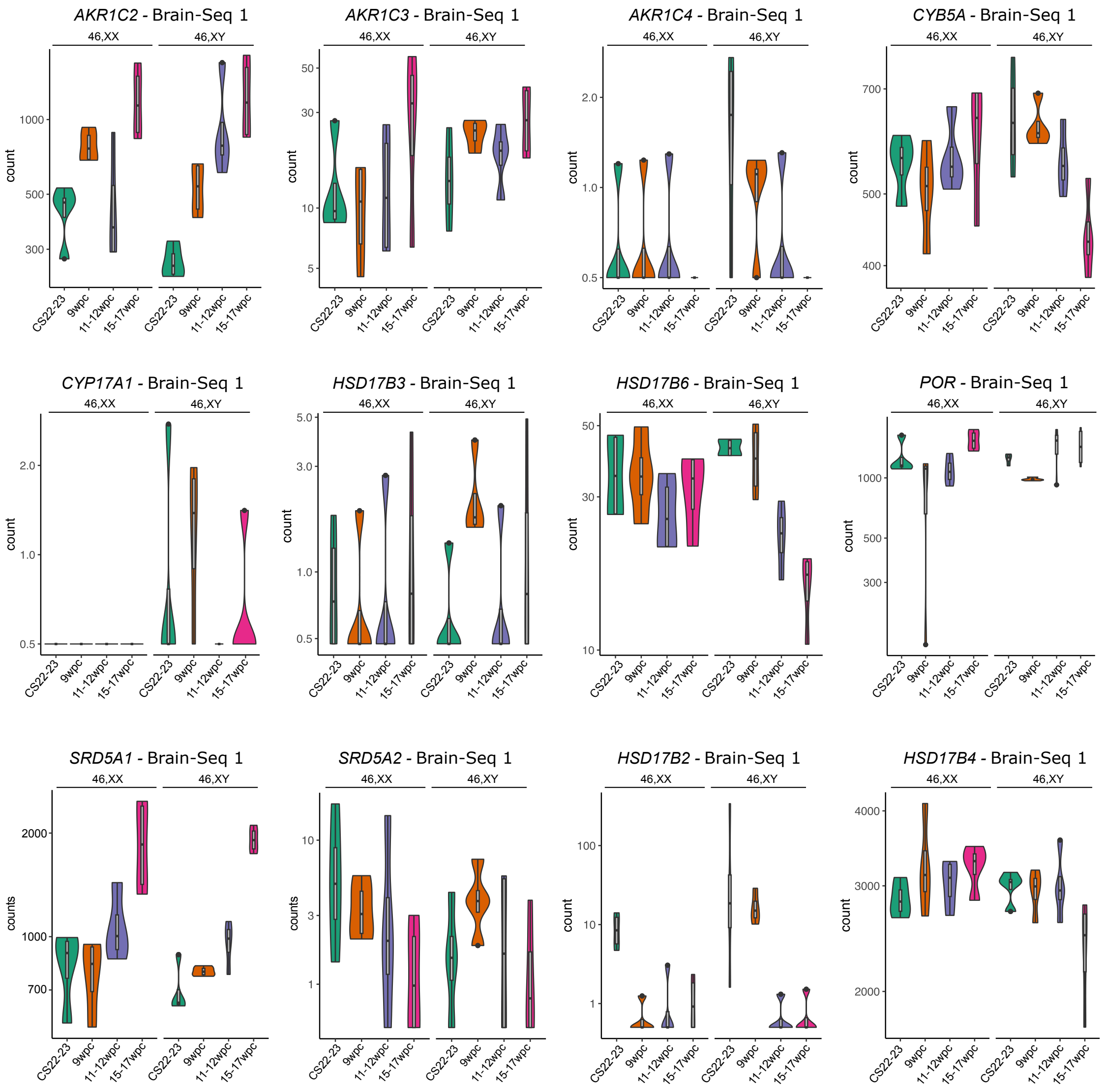

**Supplementary Figure 12. Expression of genes involved in the backdoor steroid pathway in the human brain cortex during early fetal development. a** Classic and proposed backdoor steroid biosynthesis pathway. Adapted from del Valle I et al., JCI Insight 2023;8(14):e168177; doi.org/10.1172/jci.insight.168177. **b** Data are shown as violin plots of normalized counts on a log scale, with n=4 in each group. CS, Carnegie stage; wpc, weeks post-conception.

### Supplementary Figure 13

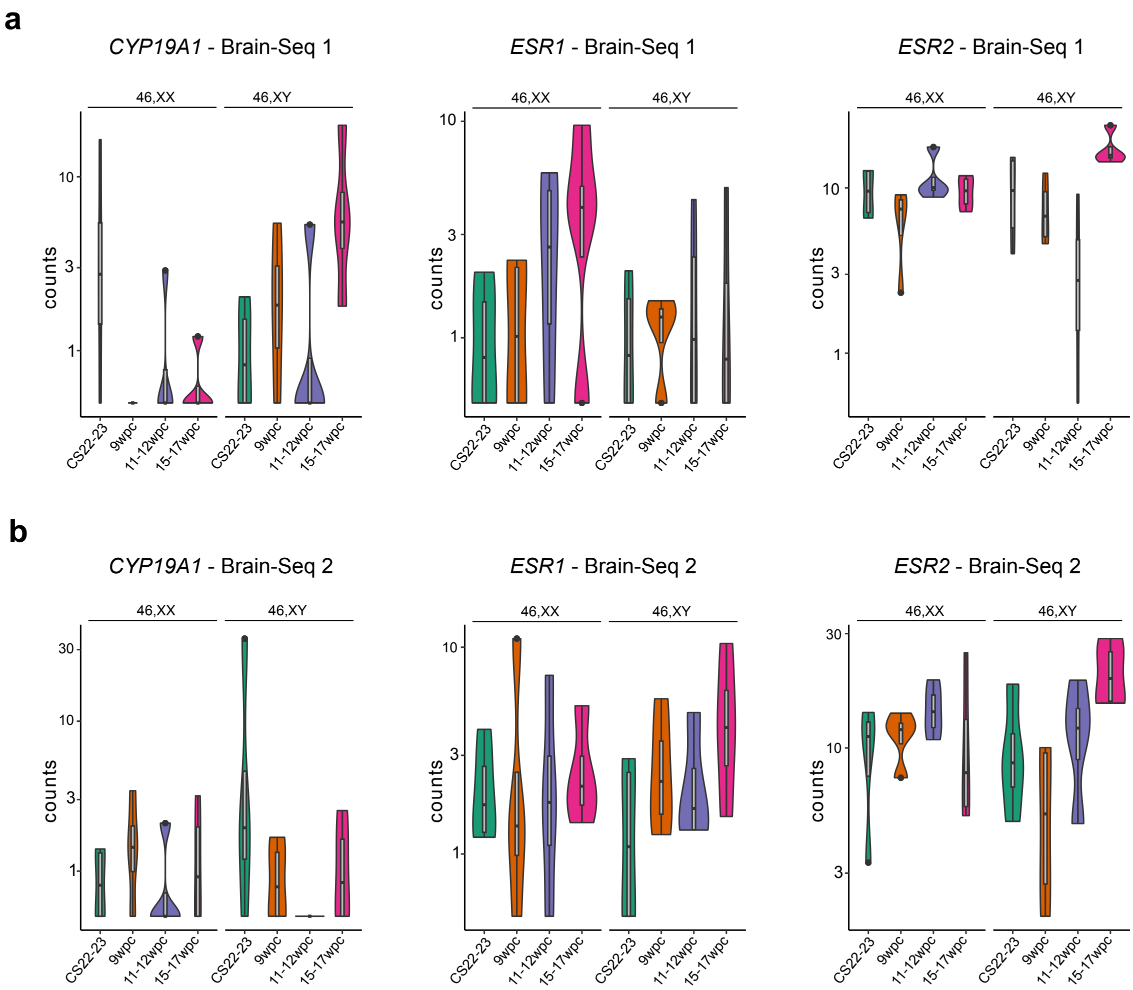


**Supplementary Figure 13. Expression of genes encoding aromatase (*CYP19A1*) and the estrogen receptors (*ESR1*, *ESR2*) in the human brain cortex during early fetal development. a** Brain-Seq 1 dataset. **b** Brain-Seq 2 dataset. Data are shown as violin plots of normalized counts on a log scale, with n=4 in each group. CS, Carnegie stage; wpc, weeks post-conception.

#### Supplementary Figure 14

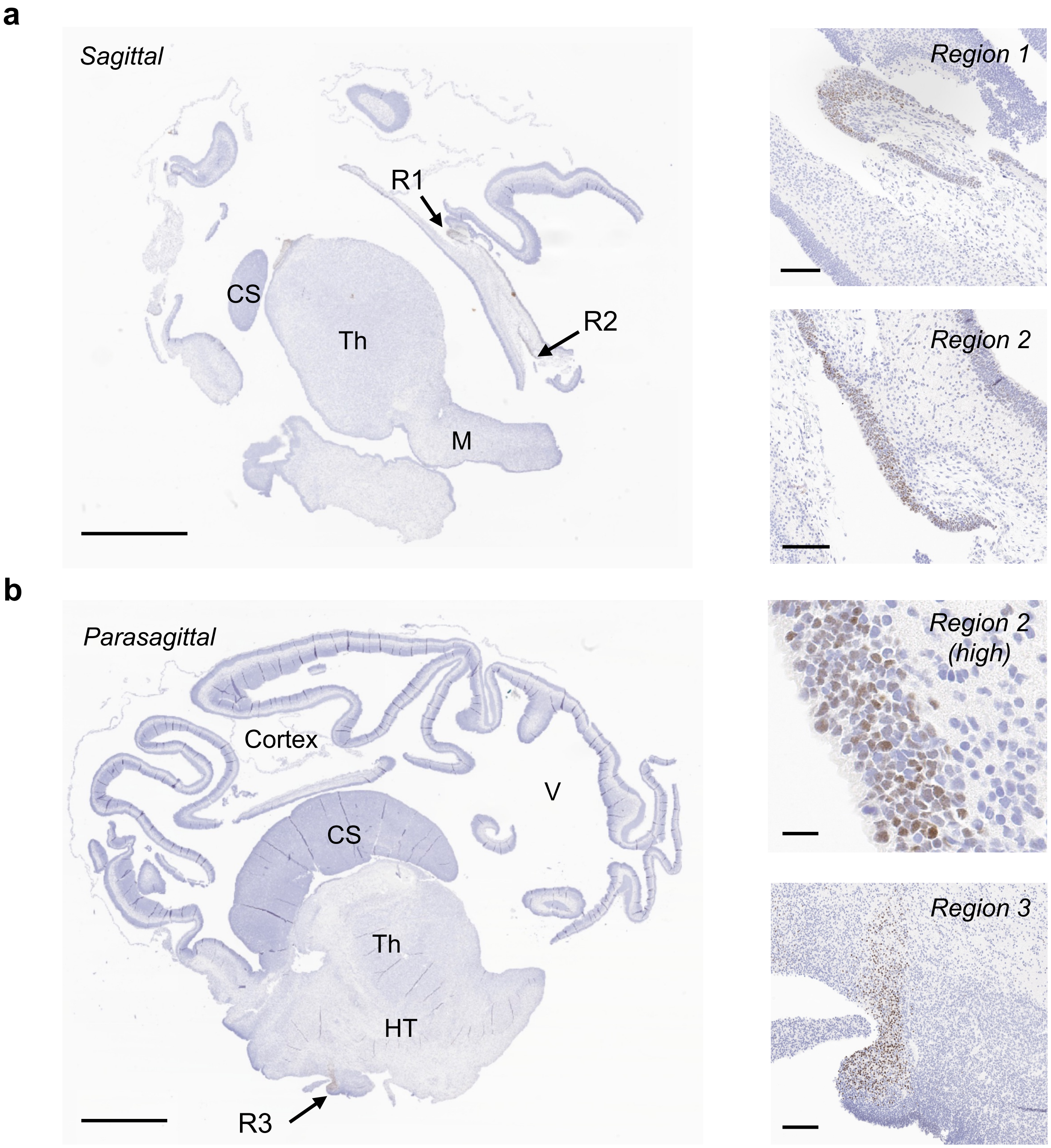


**Supplementary Figure 14. Androgen receptor expression by immunohistochemistry in the developing diencephalon and cerebral cortex at 10 weeks post conception. a** Sagittal section near the midline. **b** Parasagittal section. Arrows indicate regions of interest shown in higher magnification on the right. Sample 46,XY. Scale bars, 2mm. **c** Higher magnification images of regions of interest. Scale bars, 100 µm; Region 2 high, 20 µm. CS, corpus striatum (ganglionic eminence); HT, hypothalamus; M, midbrain; R, region; Th, thalamus.

#### Supplementary Figure 15

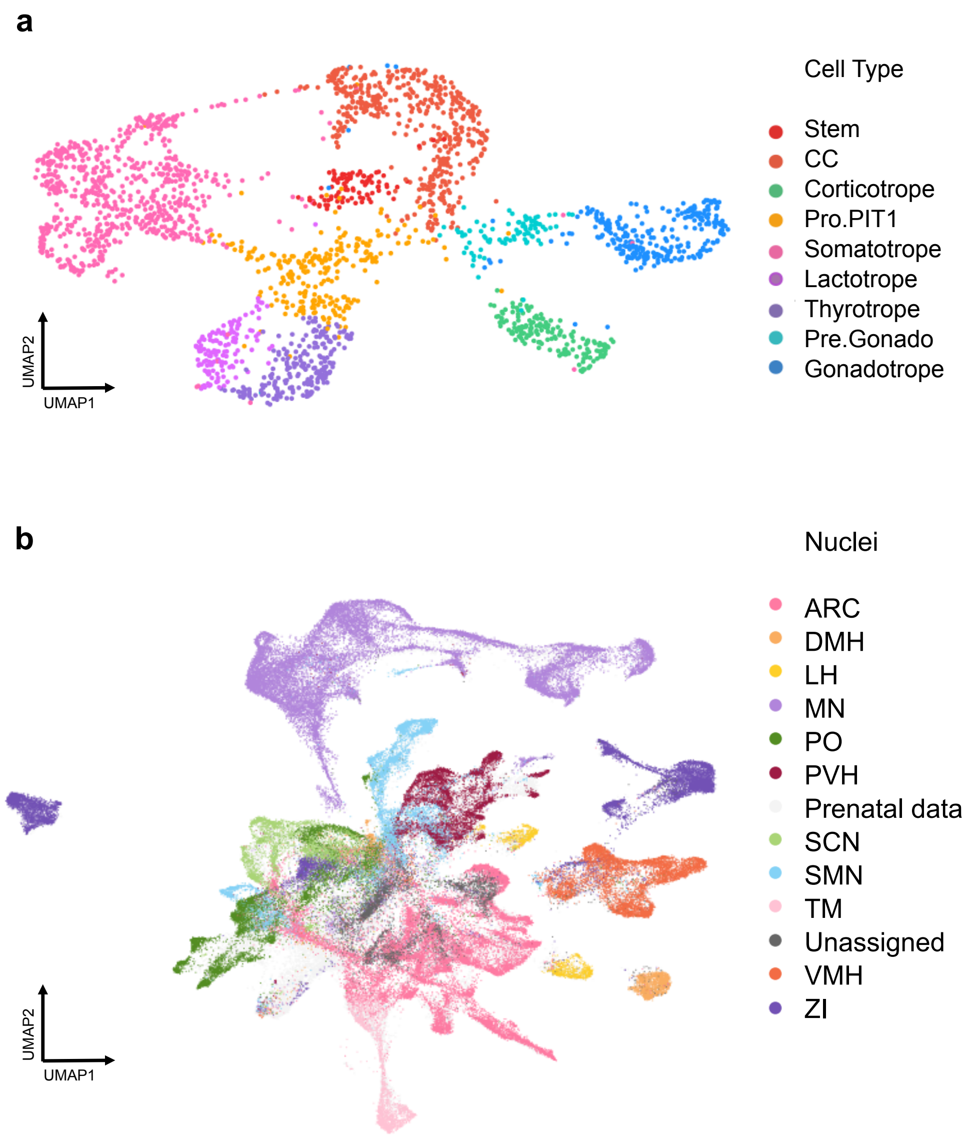


**Supplementary Figure 15. Nuclei annotations of single-cell RNA-seq data. (a)** Major cell-type clusters in the human pituitary (Zhang S et al., Nat Commun 2020. 11:5275. doi.org/10.1038/s41467-020-19012-4). CC, cell cycle cells. Pro.PIT1, progenitor cells of PIT1 lineage. Pre.Gonado, precursor cells of gonadotrope. **(b)** Nuclei identified in the human hypothalamus (Herb BR et al., Sci Adv 2023. 9,eadf6251. doi.org/10.1126/sciadv.adf6251). ARC, arcuate nucleus. DMH, dorso-medial nucleus of the hypothalamus. LH, lateral hypothalamus. MN, mammillary nucleus. PO, preoptic area. PVH, paraventricular nucleus of the hypothalamus. SCN, suprachiasmatic nucleus. SMN, supramammillary nucleus. TM, tuberomammillary nucleus. VMH, ventromedial nucleus of the hypothalamus. ZI, zona incerta. UMAP, (uniform manifold approximation and projection).
